# Maf Family Transcription Factors are Required for Nutrient Uptake in the Neonatal Gut

**DOI:** 10.1101/2022.07.26.501624

**Authors:** Anne M. Bara, Lei Chen, Celina Ma, Julie Underwood, Rebecca S. Moreci, Kaelyn Sumigray, Tongyu Sun, Yarui Diao, Michael Verzi, Terry Lechler

**Author notes:** Corresponding Author: Terry Lechler, Depts. of Dermatology and Cell Biology, Duke University Medical Center, 310 Nanaline Duke Bldg, Box 3709, Durham, NC 27710, USA.

## Abstract

There are fundamental differences in the way that neonatal and adult intestines absorb nutrients. In adults, macromolecules are efficiently broken down into simpler molecular components in the lumen of the small intestine, then absorbed. In contrast, neonates are thought to rely more on bulk intake of nutrients and subsequent degradation in the lysosome. Here, we identify the Maf family transcription factors, MafB and cMaf, as markers of terminally-differentiated intestinal enterocytes throughout life. The expression of these factors is regulated by HNF4α/γ, master regulators of the enterocyte cell fate. Loss of Maf factors results in a neonatal-specific failure to thrive and loss of bulk uptake of nutrients. RNA-Seq and CUT&RUN analyses defined an endo-lysosomal program as being downstream of these transcription factors. We demonstrate major transcriptional changes in metabolic pathways, including fatty acid oxidation and increases in peroxisome number in response to loss of Mafs. Finally, we show that deletion of Blimp1, which represses adult enterocyte genes in the neonatal gut, shows highly overlapping changes in gene expression and similar defects in nutrient uptake. This work defines transcriptional regulators that are necessary for bulk uptake in neonatal enterocytes.

## Introduction

The small intestine plays an essential role in nutrient uptake. Central to this function are enterocytes, the primary cell type responsible for absorption in the gut. Enterocytes are highly polarized columnar cells that line the villi of the small intestine. They are derived from the stem/progenitor cells that lie within the intestinal crypts and are the most abundant cell type in the intestinal epithelium. Enterocytes turnover quickly and it is estimated that billions are generated every day in adult humans (Sender and Milo, 2021).

The enterocyte cell fate is developmentally linked to the morphogenesis of the intestine. Prior to embryonic day E14.5 in the mouse, the intestine is a flat, pseudostratified epithelium (Wang et al., 2018). At E15.5, villi begin to form and enterocytes are first specified. While adult enterocytes are terminally differentiated, it has been proposed that embryonic enterocytes remain competent to contribute to the stem cell pool through fission of villi (Guiu et al., 2019).

Recent work has begun to elucidate the transcriptional pathways that control enterocyte fate. A number of transcriptional regulators, including Cdx1/2, HNF4**α/γ**, GATA4 and Elf3 play important roles in enterocyte specification (Beuling et al., 2011; Chen et al., 2019a; Chen et al., 2019b; Verzi et al., 2011). Of these, HNF4**α/γ** have emerged as essential regulators of the enterocyte program. They are required for brush border assembly as well as expression of many canonical enterocyte genes (Chen et al., 2021; Chen et al., 2019a; Chen et al., 2019b). While HNF4**α/γ** are essential for enterocyte gene expression, they are not sufficient to drive this cell fate. The expression of HNF4 **α/γ** throughout the crypt/villus axis, not exclusively in enterocytes, indicates additional factors are required to regulate enterocyte gene expression.

Our lab identified MafB as an enterocyte specific transcription factor in the intestinal epithelium (Sumigray et al., 2018). MafB and cMaf are both members of Maf family of proteins and are characterized as large Maf’s, containing both DNA binding and transactivating domains. MafB and cMaf act redundantly in both the epidermis, by regulating genes associated with differentiated keratinocytes, and in macrophages, by controlling exit from the cell cycle (Aziz et al., 2009; Lopez-Pajares et al., 2015). Here, we sought to characterize the expression of cMaf and MafB in the intestinal epithelium and determine their role in enterocytes.

Enterocytes are a heterogeneous group of cells whose gene expression is regulated both spatially and temporally. There are distinct enterocyte signatures and functions along the proximal-distal axis of the intestine, and in zones along the villus axis (Haber et al., 2017; Moor et al., 2018; Park et al., 2019). In addition, there are significant changes in enterocytes between neonatal/suckling and post-suckling/adult stages (Wilson et al., 1991). During the neonatal stage there is both rapid growth and a relative immaturity of the intestine and other organs that support digestion, such as the liver and pancreas (Greengard, 1977; Robberecht et al., 1971). The major changes in expression of digestive enzymes that occur as animals approach weaning indicate a shift in expression program to support the changes in the diet from the milk-based diet of neonates to solid food post-weaning. The differences between neonatal and adult enterocytes are reflected in fundamentally different morphologies of the apical domain of the cell (Muncan et al., 2011; Skrzypek et al., 2007), and in mechanisms of nutrient uptake. In adults, most macromolecules are broken down in the lumen of the intestine and then absorbed. In contrast, neonates are thought to rely heavily upon bulk uptake of nutrients and subsequent degradation within lysosomes (Gonnella and Neutra, 1984; Wilson et al., 1991). In support of this, we previously demonstrated that the endocytic adaptor protein, Dab2, is required for protein uptake in suckling mice (Park et al., 2019). Additionally, a recent study in Caco-2 cells determined that active, caveolae-dependent processes are required for intake of labeled pea protein; however, pharmacological inhibition of phagocytosis did not disrupt protein intake (Zhang et al., 2022). These studies have begun to identify the cellular mechanisms of nutrient uptake, however, many questions remain about the machinery required for this process, differences in uptake in neonatal and adult enterocyte, and the transcriptional regulation of these pathways.

Little is known about the transcriptional regulation of neonatal enterocyte function, with one interesting exception. The transcriptional repressor, Blimp1 (encoded by the Prdm1 gene) is expressed in neonatal mice and its expression is largely lost around weaning. Ablation of the Prdm1 gene in the intestinal epithelium results in premature expression of adult enterocyte gene signature (Harper et al., 2011; Muncan et al., 2011). However, it is not known whether Blimp1 is required for bulk protein uptake in the neonatal intestine in addition to its role in repressing adult genes.

Here, we demonstrate that the transcription factors MafB and cMaf are expressed in enterocytes beginning at the earliest stage of their embryonic specification and through adulthood. Their expression is controlled by the enterocyte master regulators, HNF4α/γ. Further, we show that loss of Maf factors results in a failure to thrive and an inability of neonatal enterocytes to take up nutrients in bulk from the intestinal lumen. In addition, we find that loss of Blimp1 results in very similar changes in both gene expression and in an inability to take up nutrients. Together these data begin to define the transcriptional regulation of nutrient uptake in the neonatal gut.

## Results

### Developmental Expression of Mafs

Enterocytes are first specified around embryonic day 15.5 (E15.5) of mouse development, concomitant with the folding of villi. Prior to this, the intestinal epithelium is a flat pseudostratified epithelium (Wang et al., 2018). Using two different antibodies – one that recognizes both MafB and cMaf (Mafs), as well as one that uniquely recognizes cMaf (Figure 1, Supplement 1), we did not detect protein expression of Maf factors in the E14.5 intestinal epithelium. A day later, at E15.5, Maf factors were found in the nuclei of cells at the tops of the emerging villi (Figure 1A). At both stages, there were also occasional mesenchymal cells that were Maf positive. To determine the relationship between villi folding and Maf expression, we quantitated villi height and the emergence of Maf expression. There was a clear demarcation in villus height and Maf expression as emerging villi shorter than 50μm were largely Maf negative, while almost all villi taller than 50μm contained Maf-expressing cells (Figure 1B).

**Figure 1.**
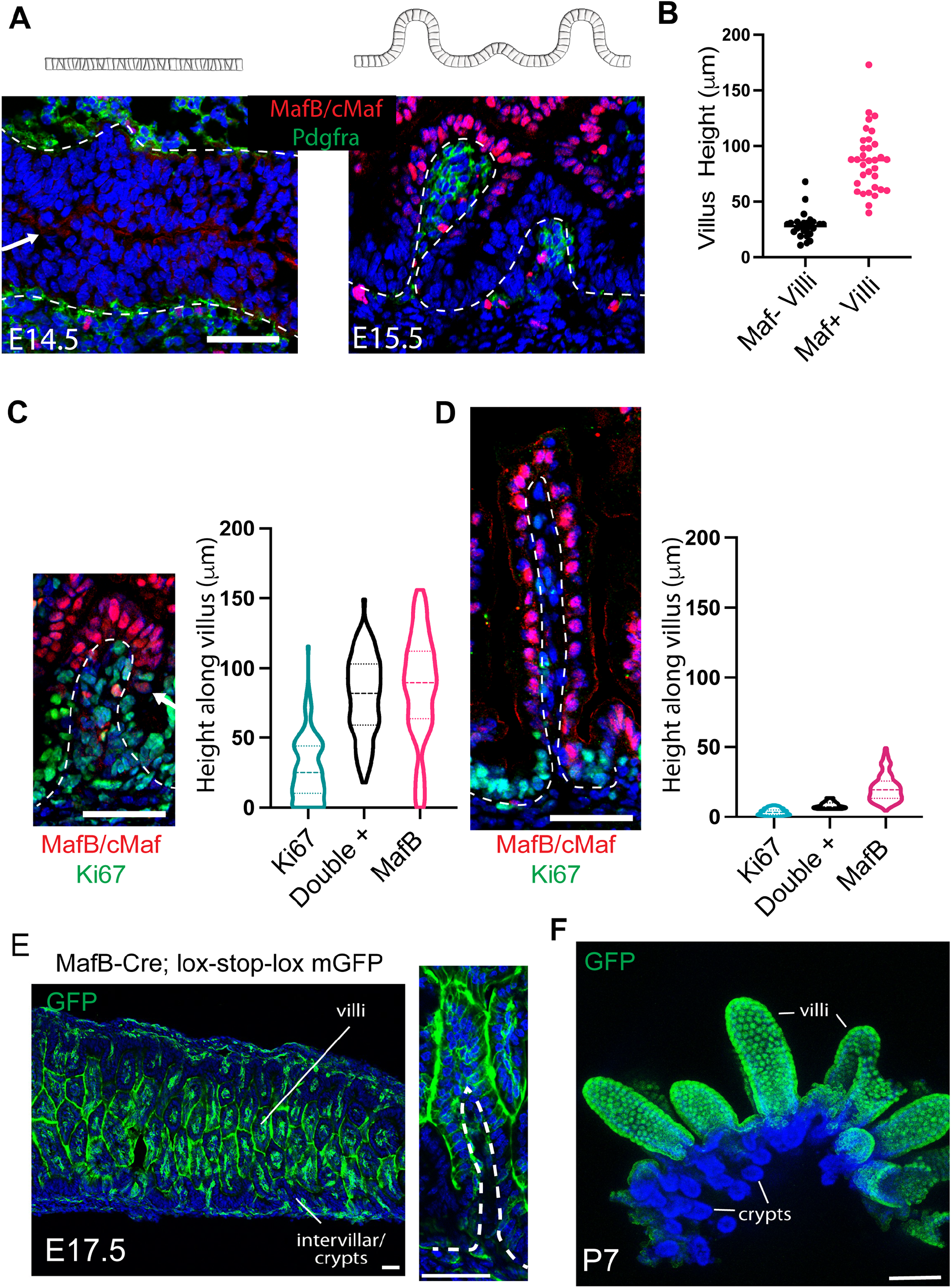
Maf proteins are expressed from the onset of enterocyte differentiation. (A and B)Expression of MafB (red) in the developing intestine at E14.5 before villi emerge (A) and at E15.5, once villi have begun to form (B). Basement membranes are outlined in white; the arrow indicates the epithelium. Cartoon depicts intestinal morphology at indicated developmental stage. (B) Quantification of Maf expression by villus height at E15.5. Villus height was measured from the base of the villus to the PDGFRa cluster at the top of the villus. Villi were grouped in the 2 categories – either expressing Maf proteins (n=33), or negative for Maf proteins (n=25). (C and D) Quantification of Maf protein (red) and Ki67 (green) colocalization at E15.5 (C) and P0 (D) The positions of individual nuclei were measured by their height along the villus axis. Each cells was classified as either Ki67+ only (n=166 cells from 3 mice at E15.5, n= 135 at P0), Maf+ only (n=102 at E15.5, n= 323 at P0), or Ki67/Maf double + (n=102 at E15.5, n=11 at P0). All nuclei over 50μm from the villus base were Maf+ at P0. (E and F) Lineage trace of MafB + cells. MafB-Cre driving the expression of GFP. By E17.5 (E) a majority of embryonic enterocytes are labeled with GFP. The image on the right shows a villus at higher magnification. No MafB labeled cells are detected in crypts of P7 mice (F).

Since differentiation is typically associated with loss of proliferation, we sought to determine whether there is a correlation between the expression of Maf proteins and loss of proliferation. At E15.5, the Maf negative cells in inter-villar spaces and cells less than 50μm from the base of villi tended to express the proliferation marker, Ki67. Conversely, cells over 50μm from the base of villi generally expressed Maf proteins and many cells at this stage (27.5%) were positive for both Ki67 and Mafs (Figure 1C). At postnatal day 0 (P0), the day the mice are born, the Ki67+ region was restricted to inter-villar spaces and the 10μm at the base of villi. Epithelial cells over 10μm from the base of the villi were negative for Ki67 and most express Maf proteins (Figure 1D). Together, these data demonstrate that Maf factors are expressed at the earliest stages of enterocyte specification and that cells become post-mitotic shortly after Maf expression.

To verify these results, we took advantage of an existing MafB-Cre knock-in mouse line and a fluorescent reporter to visualize and lineage trace MafB expressing cells (Muzumdar et al., 2007; Wu et al., 2016). We observed MafB lineage-labeled cells in the embryonic intestine, with a pattern similar to endogenous Maf expressing cells, but shifted slightly later in developmental time, likely reflecting the delay in recombination and subsequent expression of the GFP reporter. At E16.5, 24% of epithelial cells were fluorescently labeled as determined by FACS sorting (7 mice, >1000 cells counted/mouse) and by E17.5 most cells on villi were lineage labeled (Figure 1E). In postnatal mice, villi were uniformly labeled while crypts were negative, fewer than 1% of crypts were labeled at P7 (Figure 1F). While these data are consistent with our antibody staining, they are inconsistent with a proposed model that embryonic villi undergo fission with villar cells giving rise to adult stem cells (Guiu et al., 2019). Our findings demonstrate that MafB positive embryonic enterocytes do not substantially contribute to the adult stem cell lineage. These data are incongruent with the idea that all embryonic intestinal epithelial cells have equal abilities to contribute to adult stem cells (Guiu et al., 2019). Importantly, the modeling studies of K19-Cre^ER^ lineage traced cells assumed lineage labeling occurred at equal frequencies in villar and inter-villar cells (Guiu et al., 2019). However, when we analyzed recombination 24 hours after K19-Cre^ER^ induction, we found that there was a preference for recombination to occur within the inter-villar regions (Figure 1, Supplement 2). This may reflect differences in expression of K19 in different cell types, or it may reflect increased recombination activity in proliferative cells, which has been previously noted (Mascre et al., 2012). In either case, our data demonstrate that not all embryonic intestinal cells give rise to adult stem cells and the MafB positive population is a differentiated cell type that does not contribute to adult stem/crypt cells.

### Maf factor expression is regulated by HNF4α/γ

As Maf factors mark differentiated enterocytes, we wanted to address the upstream transcriptional regulators of their expression. HNF4**α/γ** transcription factors are crucial for enterocyte specification and intestinal homeostasis (Chen et al., 2021; Chen et al., 2019a; Chen et al., 2019b). We therefore analyzed mice in which HNF4**α/γ** were deleted throughout the intestinal epithelium, using Villin-Cre^ER^ (el Marjou et al., 2004). Adult mice were collected 3 days after deletion of HNF4**α/**γ was initiated in the intestinal epithelium (HNF4 DKO). Multi-omic analysis of these mice has previously been performed, and we mined those datasets to determine effects of HNF4 DKO on Maf factor expression (Figure 2 A,D,E) (Chen et al., 2019a; Chen et al., 2019b). RNA-Seq analysis demonstrated a substantial reduction in both MafB and cMaf mRNA in HNF4 DKO intestinal epithelia (Figure 2A). This result was validated at the protein level – with a decrease in both the intensity of staining and the percentage of MafB positive cells in the mutant tissue (Figure 2 BC). In addition, we found that HNF4**α/γ** binding was enriched on/around the Maf and MafB genes by ChIP-Seq analysis (Figure 2D), suggesting that this regulation may be direct. Further, the content of H3K27ac was clearly reduced at these loci in HNF4 DKO mice, consistent with a conversion to less active transcription (Figure 2D). Finally, we examined chromatin looping in control and HNF4 DKO intestines. The interactions between chromosomal domains were clearly decreased at the MafB and cMaf loci in HNF4 DKO intestine (Figure 2E). Together, these data indicate that Maf factors are direct transcriptional targets of HNF4**α/γ**.

**Figure 2.**
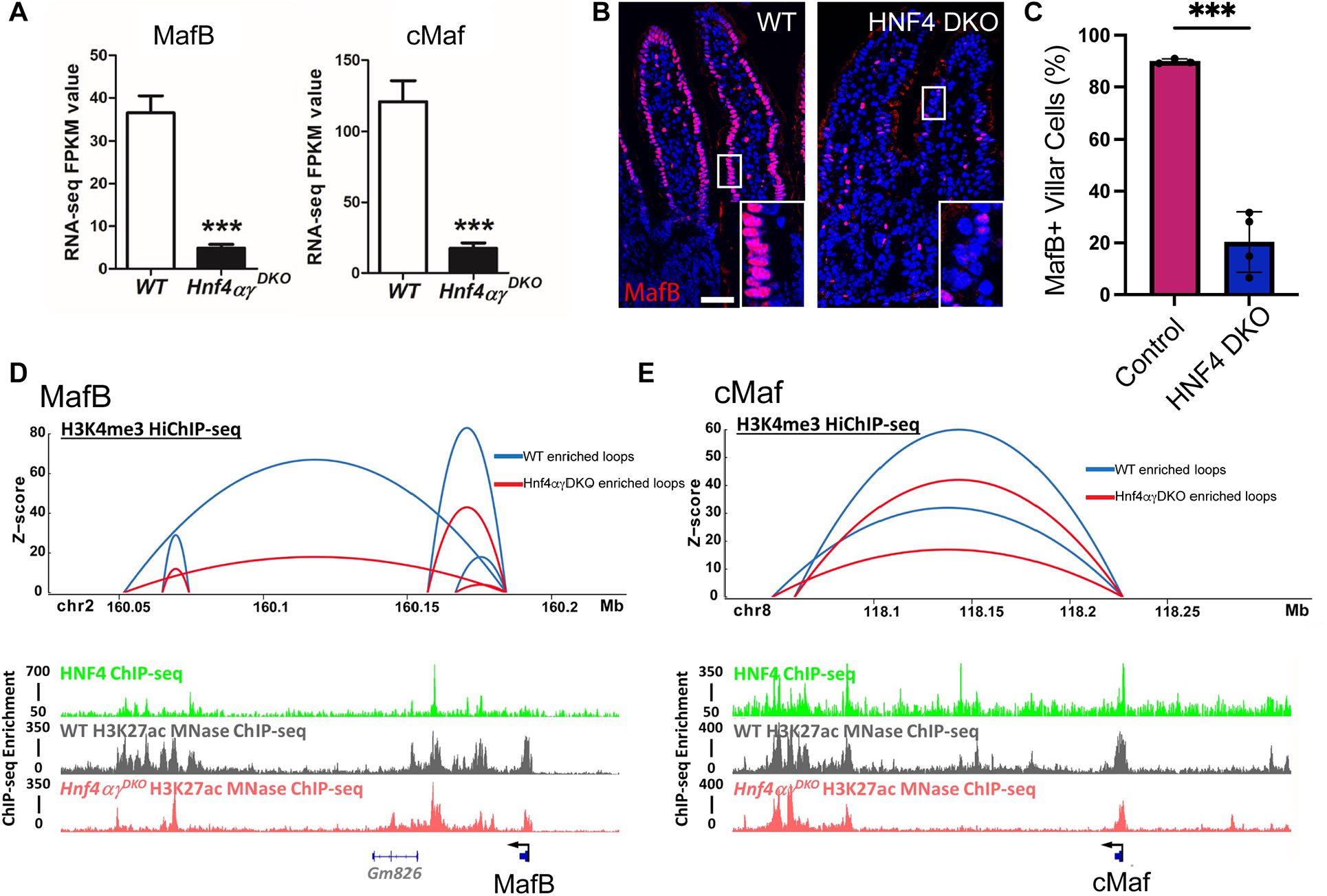
Maf proteins are downstream of HNF4 factors in the intestinal epithelium. (A) Analysis of MafB and cMaf mRNA levels by RNA sequencing of HNF4 DKO adult mice 3 days after tamoxifen administration. (B) Immunostaining for Maf proteins (red) in control and HNF4 DKO tissue. (C) Quantification of the number of Maf positive nuclei on villi of control and HFN4 DKO intestines (n=3 control, 2835 cells counted and 4 HNF4 DKO mice, 4036 cells counted. (D and E) MafB (D) and cMaf (E) genetic loci are depicted. Top images show DNA looping in control (blue) and HNF4α/γ DKO (red) intestines at gene loci of MafB and cMaf. Peaks for HNF4α/γ binding on MafB and cMaf genes (Green), WT H3K27ac (grey), and HNF4 DKO H3K27ac (orange). Arrow indicated transcriptional start site.

### Loss of Maf factors results in a failure to thrive

To examine the function of Maf proteins in the intestinal epithelium, we obtained floxed alleles for MafB and cMaf and used Villin-Cre and Villin-Cre^ER^ to drive their conditional deletion in the intestinal epithelium (Maf DKO and Maf iDKO respectively) (Wende et al., 2012; Yu et al., 2013). Villin-Cre is active in the intestinal epithelium starting at E14.5; therefore, in this model, the intestinal epithelium never expresses Maf factors (Figure 3A). Immunofluorescence analysis confirmed the loss of Maf proteins in Maf DKO mice, demonstrating efficient deletion, yet there were no major changes in the morphology of the intestine (Figure 3B). Maf DKO mice were born at Mendelian ratios; however, mutant mice were clearly identifiable as being smaller than littermate controls by P3. From this time onward, Maf DKO mice were about 70% the weight of control mice, a phenotype consistent with deficits in intestinal function (Figure 3 C,D). While some animals needed to be euthanized at weaning because of their small size, most survived into adulthood and were able to reproduce, though their smaller size persisted.

**Figure 3.**
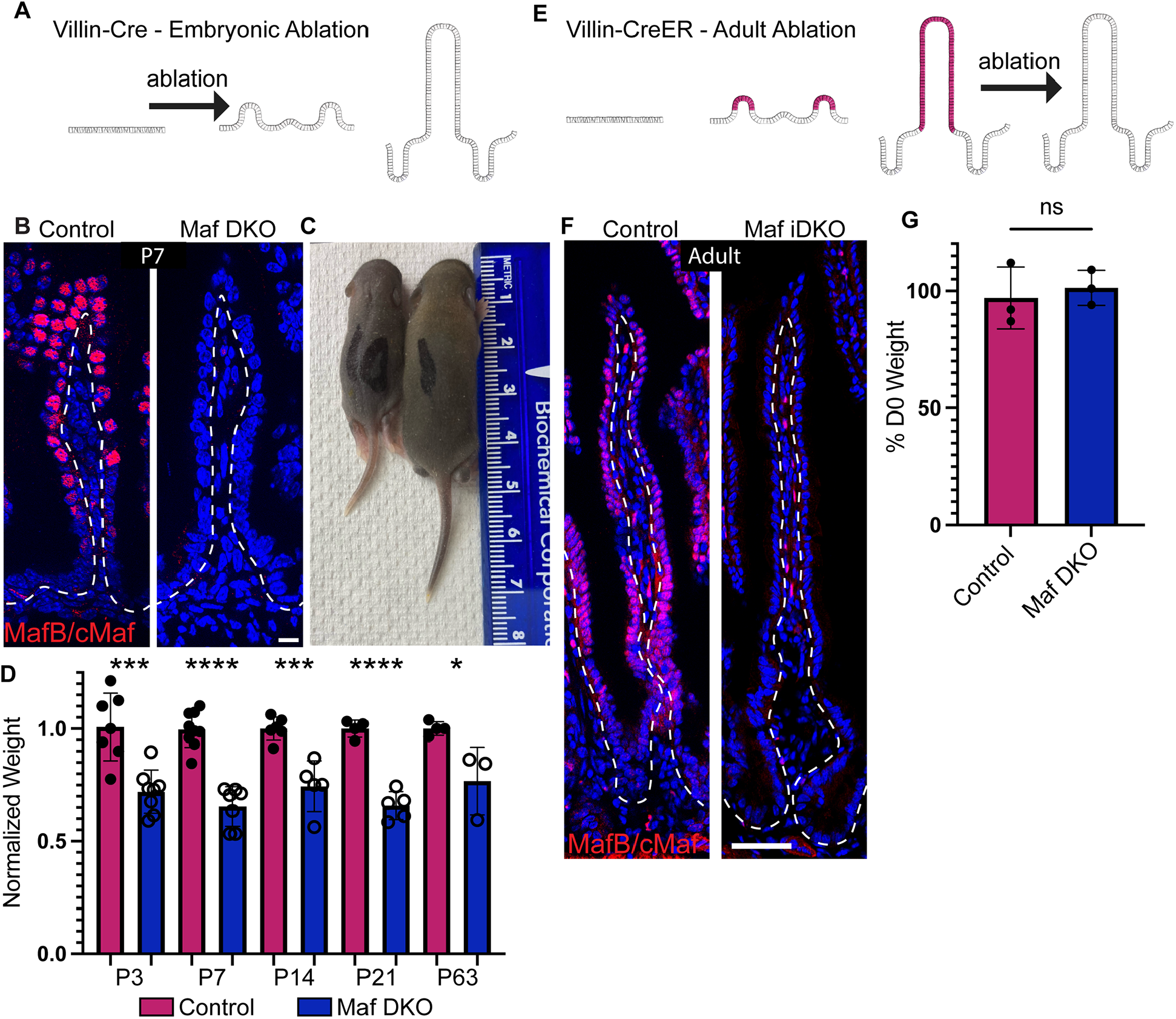
Developmental loss of Maf proteins leads to decreased body weight of neonates. (A to D) Embryonic ablation of MafB/cMaf, driven by VillinCre. (A) Schematic showing intestinal epithelium developing without Maf proteins (red). (B) Control and Maf DKO villus at P7, Maf proteins (red), basement membrane outlined in white, scale bar 10 μm. (C) Maf DKO (Left) and control (right) mice at P7. (D) Weight of Maf DKO mice as a percent of control littermate average weight. (E) Schematic showing normal Maf expression during development and deletion of MafB/cMaf in adult mice after addition of tamoxifen. (F) Control villus and Maf DKO villus in adult mice. Maf proteins (red), basement membrane outlined in white, scale bar 25μm. (G) Weights of control and Maf iDKO mice 30 days after the first of 2, 2.5mg IP tamoxifen injections at D0 and D2 as a percent of their initial body weight. n=3 control, n=3 Maf DKO Unpaired T-test was used to compare the samples, p=0.6475.

To determine potential functions of Maf proteins in adult mice, we used the Villin-Cre^ER^ driver and tamoxifen to delete MafB and cMaf after weaning (Maf iDKO) (Figure 3E). Similar to the developmental deletion of Maf proteins, no major changes in morphology were detected in Maf iDKO mice (Figure 3F). Weights of Maf iDKO mice and controls were measured weekly for 1 month following tamoxifen injection. The weights of Maf iDKO mice did not differ from controls (Figure 3G). Therefore, Maf factors play a clear role in intestinal function at neonatal stages, but are largely dispensable for intestinal morphology and function in adulthood. Further analysis is needed to determine whether specific pathways are disrupted upon Maf DKO in adults.

Consistent with the normal epithelial architecture, we did not detect changes in proliferation following deletion of MafB and cMaf whether the deletion occurred during development or postnatally (Figure 3, Supplement 1A,B, E,F, I,J). We also asked whether loss of enterocyte specific factors leads to a mis-regulation of lineage specification and results in increased numbers of secretory cells. We quantified the percent of secretory goblet cells using Muc2 as a marker and observed no change in Maf DKO neonates or adults or Maf iDKO adults (Figure 3, Supplement 1CD, GH, KL). Therefore, while decreased body weight of Maf DKO mice suggests functional consequences of Maf loss during development, no major changes were detected in gross morphology, proliferation, and secretory lineage specification.

### Maf factors are required for expression of phagocytic and endo-lysosomal genes

In order to assess the transcriptional changes resulting from Maf DKO, we turned to bulk RNA sequencing. We isolated epithelial cells from proximal, medial, and distal intestine of 3 control and 3 Maf DKO mice at postnatal day 7 and pooled all 3 regions into a single tube per mouse. Bulk RNA sequencing indicated that over 3500 genes were differentially expressed in Maf DKO mice (log2 fold change>0.25, and p<0.05), with similar numbers of up- and down-regulated genes (Figure 4 A,B). Maf proteins are thought to function largely as transcriptional activators, thus we initially focused on genes that were down-regulated in the mutant. KEGG pathway analysis of this gene set revealed a striking down-regulation of genes involved in lysosomes, endocytosis, and phagocytosis (Figure 4C). These are all pathways implicated in the bulk uptake and degradation of nutrients. In adults, enterocytes express a diverse array of digestive enzymes and nutrient transporters which enable initial nutrient breakdown to occur extracellularly, and digested nutrients are transported into the enterocytes. In neonates, these pathways are not yet mature and enterocytes are thought to rely on bulk intake of nutrients and subsequent digestion in lysosomes (Gonnella and Neutra, 1984; Wilson et al., 1991). However, much of the machinery regulating this intake, as well as the transcriptional regulation of neonatal nutrient uptake, has not been identified.

**Figure 4.**
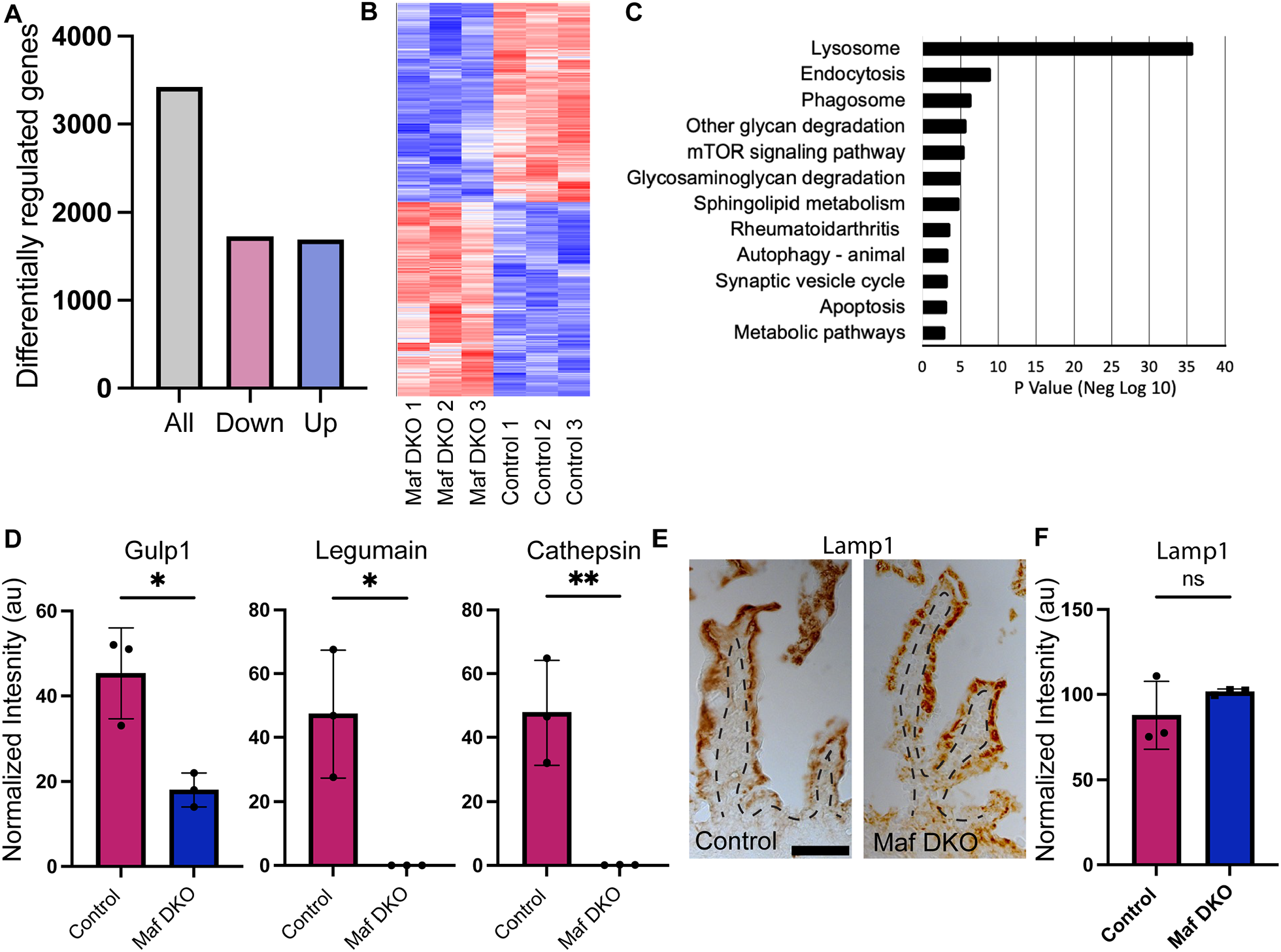
Maf proteins control expression of endosomal, phagocytic and lysosomal genes in the neonatal intestine. (A and B) Epithelial cells were isolated from intestines of P7 mice n=3 control and n=3 Maf DKO mice. Cells were collected from proximal, medial, and distal regions and pooled into a single sample for each mouse. Bulk RNA sequencing identified 3425 differentially regulated genes, 1731 down regulated and 1694 upregulated (p<0.05). (C) KEGG analysis of down regulated genes. (D) Western blots of protein isolated from distal intestinal epithelium of P7 mice. n=3 control and n=3 Maf DKO showing decreased protein expression of Gulp1 (p=0.0143), Legumain (p= 0.0147), and Cathepsin (p=0.0073). Unpaired T-Test. Band intensities were normalized to tubulin loading control. (E) Immunohistochemistry stain for Lamp1 in P7 intestinal epithelium of control and Maf DKO (scale bar 50 μm). (F) Western blot analysis of Lamp1 levels.

To validate this transcriptomic data, we examined the levels of proteins predicted to be down-regulated in the Maf DKO. The phagocytic receptor, GULP1, as well as legumain and cathepsins (both lysosomal proteases) were all detected at lower levels in intestinal epithelial lysates collected from the Maf DKO consistent with the RNA-Seq results (Figure 4D). Given the decrease in expression of many lysosomal genes and proteins, we assessed lysosomal architecture in the Maf DKO enterocytes. While many of the mRNAs for lysosomal enzymes were dramatically decreased, those encoding structural components were only modestly decreased (Lamp1 down by 1.72 fold and Lamp2 by 1.66 fold). Consistent with this, we did not see a significant decrease in lysosomal number as assayed by Lamp1 staining or protein level via western blot (Figure 4 E,F). This suggests that the function but not the biogenesis of lysosomes is affected in Maf DKO intestine.

We next sought to identify chromosomal loci bound by MafB using CUT&RUN analysis. Two independent samples were generated using cells isolated from P7 villi. We identified ∼4000 bound loci that were present in both samples, roughly equivalent to the number of loci bound by MafB in keratinocytes (Lopez-Pajares et al., 2015). Importantly, comparison to the genes identified as both down regulated in Maf DKO and loci bound by MafB revealed KEGG terms for endocytosis, phagocytosis and lysosomes, supporting the idea that Maf factors may directly regulate these targets (Figure 4, Supplement 1). Analysis of genes both down-regulated in Maf DKO intestines and bound by MafB revealed a statistically significant enrichment of shared genes (p<4.0×10^-41^), thus supporting regulatory functions of MafB at these loci. When analyzing genes that are both bound and down-regulated by Maf factors, actin regulators was the top-enriched term. Of the 13 actin regulators on this list, 11 have been demonstrated to play roles in endocytosis or phagocytosis. Further, Gulp1 (a phagocytic receptor), Dab2 (an endocytic adapter), Lrp2 (a scavenger receptor), and the endocytic recycling proteins Rab11a and Rab11fip2 were identified as bound by MafB (Figure 4, Supplement 1). Together, these data support a model in which Maf proteins directly regulate the expression of genes necessary for bulk intake of nutrients.

### Neonatal Maf cDKO enterocytes fail to take up protein and dextran

The decreased expression of genes/proteins important for endocytosis and lysosome function suggests that these mice might have defects in nutrient uptake and degradation. In suckling neonates, enterocytes internalize milk which contains components that are auto-fluorescent, especially in the green wavelengths. The intake of auto-fluorescent compounds from the milk is especially evident in the distal intestine (Figure 5A). However, this autofluorescence was not observed in the intestines of Maf DKO neonates (Figure 5B). The absence of milk fluorescence in Maf DKO enterocytes was not secondary to lack of feeding as both control and Maf DKO pups had milk in their stomachs and throughout the lumen of the intestine. These observations demonstrate that failure to take in nutrients is at the level of the enterocytes.

**Figure 5.**
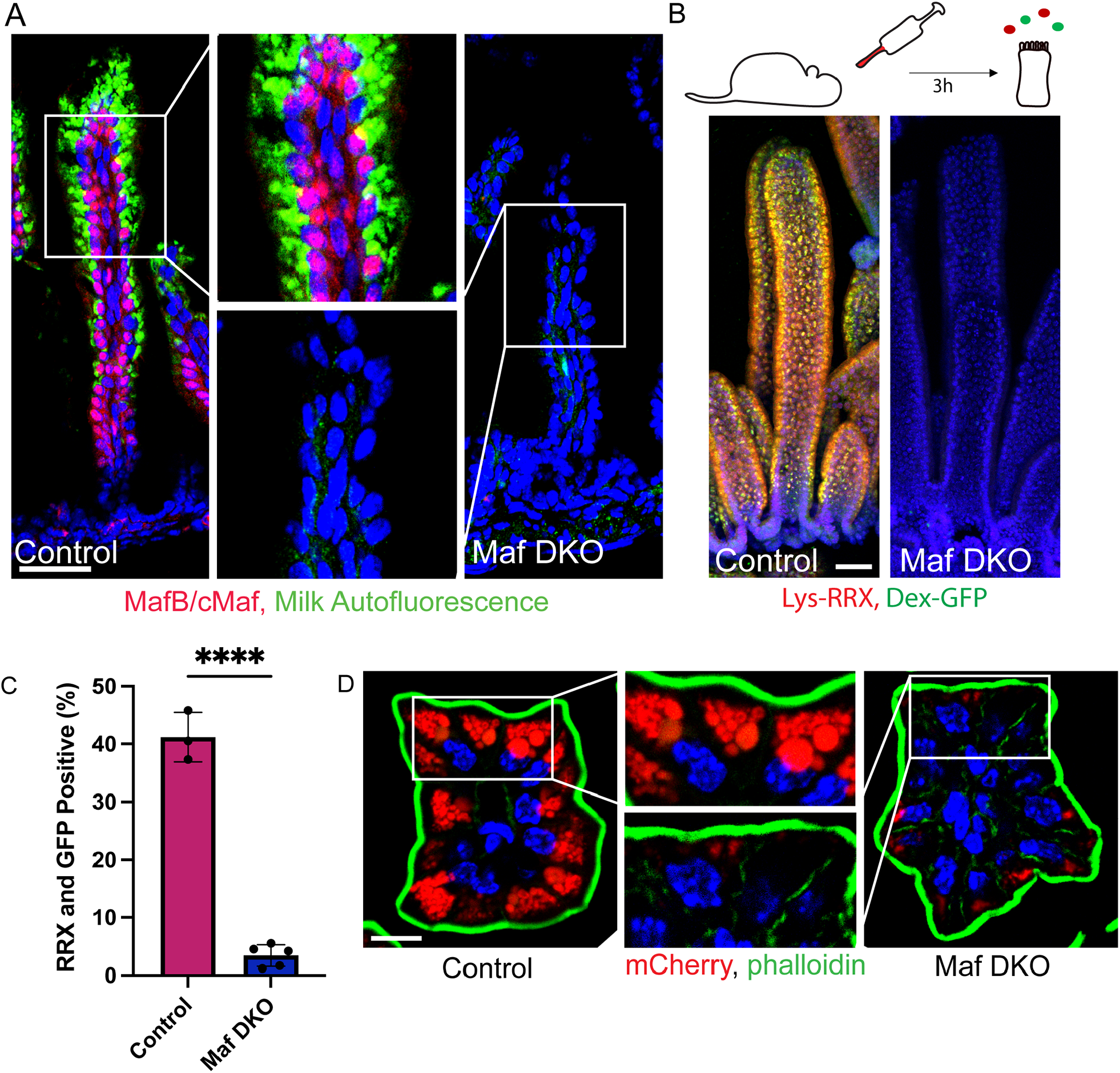
Defective bulk intake of nutrients in Maf DKO enterocytes. (A). Autofluorescence from milk (green) taken in by control enterocytes is not detected in Maf DKO mice. Images from the distal intestine of P7 mice. MafB is in red. Scale bar 50 μm. (B) (Top) Schematic for experimental set up. P7 control and Maf DKO mice were gavaged with 40 μg fluorescently labeled protein (Lysozyme-RRX, red) and 40 μg fluorescently labeled carbohydrate (Dextran-GFP, green). After 3h, mice were sacrificed and intestines were collected. (Bottom) Maximum intensity projections of intestinal epithelium collected from the medial intestine of control and Maf DKO mice are shown. Scale bar 50 μm. (C) Quantification of uptake in individual cells via FACS sorting. n= 3 control, and n=5 Maf DKO. T-test was used to compare the samples, p=<0.0001. 10,000 cells/sample were counted. (D) Cross sections of villi from mice gavaged with mCherry protein (red) 3h prior to collection. Phalloidin was used to stain actin (green). mCherry protein is found in many vesical structures in enterotypes of control mice but is absent or decreased in Maf DKO enterocytes. Scale bar 10 μm.

To further probe the ability of Maf DKO enterocytes to take in nutrients in a more controlled manner and to determine if the uptake of specific classes of nutrients is affected, we turned to a gavage assay. In this assay we used oral gavage to deliver fluorescently labeled nutrients directly to the stomachs of P7 Maf DKO and control mice. To assess protein uptake, we used lysozyme tagged with rhodamine red (Lys-RRX). In addition, we used a dextran conjugated to 488 (Dex-488) as a carbohydrate macromolecule that can be taken up via bulk endocytic pathways. Three hours after oral gavage, we collected the intestines (Figure 5C). First, we examined the fluorescent content in intact intestines and observed green and red signal throughout the lumen of the small intestine and colon in both control and Maf DKO mice. Therefore, the gavaged solution had progressed throughout the digestive tract. Next, we used fluorescence microscopy to determine whether uptake of the Lys-RRX and Dex-488 could be detected in enterocytes. Epithelial whole mounts of control mice show efficient uptake of both lysozyme and dextran; however, very little uptake of labeled nutrients was detected in enterocytes of Maf DKO mice (Figure 5C). These differences were quantified using fluorescence flow sorting which revealed strongly diminished uptake capacity of Maf DKO enterocytes (Figure 5D). Closer examination of control enterocytes in cross section showed several discrete compartments of varying sizes filled with labeled protein. Smaller vesicles were located near the apical region of the cells, and larger structures, presumable lysosomes, were seen in some cells closer to the nucleus (Figure 5D). Taken together, the decreased expression of phagocytic, endocytic and lysosomal genes and resulting inability to take in protein and dextran demonstrates that Maf proteins are crucial for bulk intake in neonatal enterocytes.

### Metabolic changes in Maf DKO enterocytes

While there was a dramatic loss of bulk uptake of proteins and carbohydrates, Maf DKO mice are still able to grow, albeit at a reduced rate. To determine whether there were other transcriptional changes that may promote growth, we examined the genes that were upregulated following loss of Maf proteins. KEGG analysis indicated major rewiring of metabolic pathways. Interestingly, genes required for fatty acid degradation and peroxisomes, organelles specialized for lipid break down (Lazarow and De Duve, 1976), were upregulated (Figure 6A). In addition, genes for fatty acid transfer proteins (Slc27a4/a2), microsomal triglyceride transfer proteins (MTTP) and even apolipoproteins and their regulators (Apoc1/2/3, Apobec1, Apoa1, Apoe, Apoa4) were increased. This upregulation suggests increased lipid metabolism in Maf DKO mice. To interrogate this possibility, we examined lipid uptake in Maf DKO neonates. We gavaged P7 mice with corn oil and collected the intestines 3 hours later. We used Oil Red O to stain the lipids and observed that both control and Maf DKO mice efficiently take in lipids (Figure 6B, C). The ability of Maf DKO enterocytes to take in lipids despite down regulation of machinery for bulk in take is consistent with the fact that fatty acids and cholesterol do not rely on bulk intake pathways (Johnston and Borgstroem, 1964; Shiau, 1981). Next, we stained for the peroxisomal marker, Pmp70, to determine whether the increased levels of peroxisomal mRNAs reflected changes in the levels of this organelle. We found a clear increase in Pmp70-stained peroxisomes (Figure 6 D,E). These data, in addition to the up-regulation of fatty acid β-oxidation enzymes (Figure 6A), point to potential compensation for lack of protein and carbohydrate intake through increased metabolism of lipids. Further, the expression of enzymes for the biosynthesis of amino acids and cofactors was also up-regulated (Figure 6A). This could indicate use of metabolic building blocks from lipid break down to synthesize other metabolites that are limited in Maf DKO mice.

**Figure 6.**
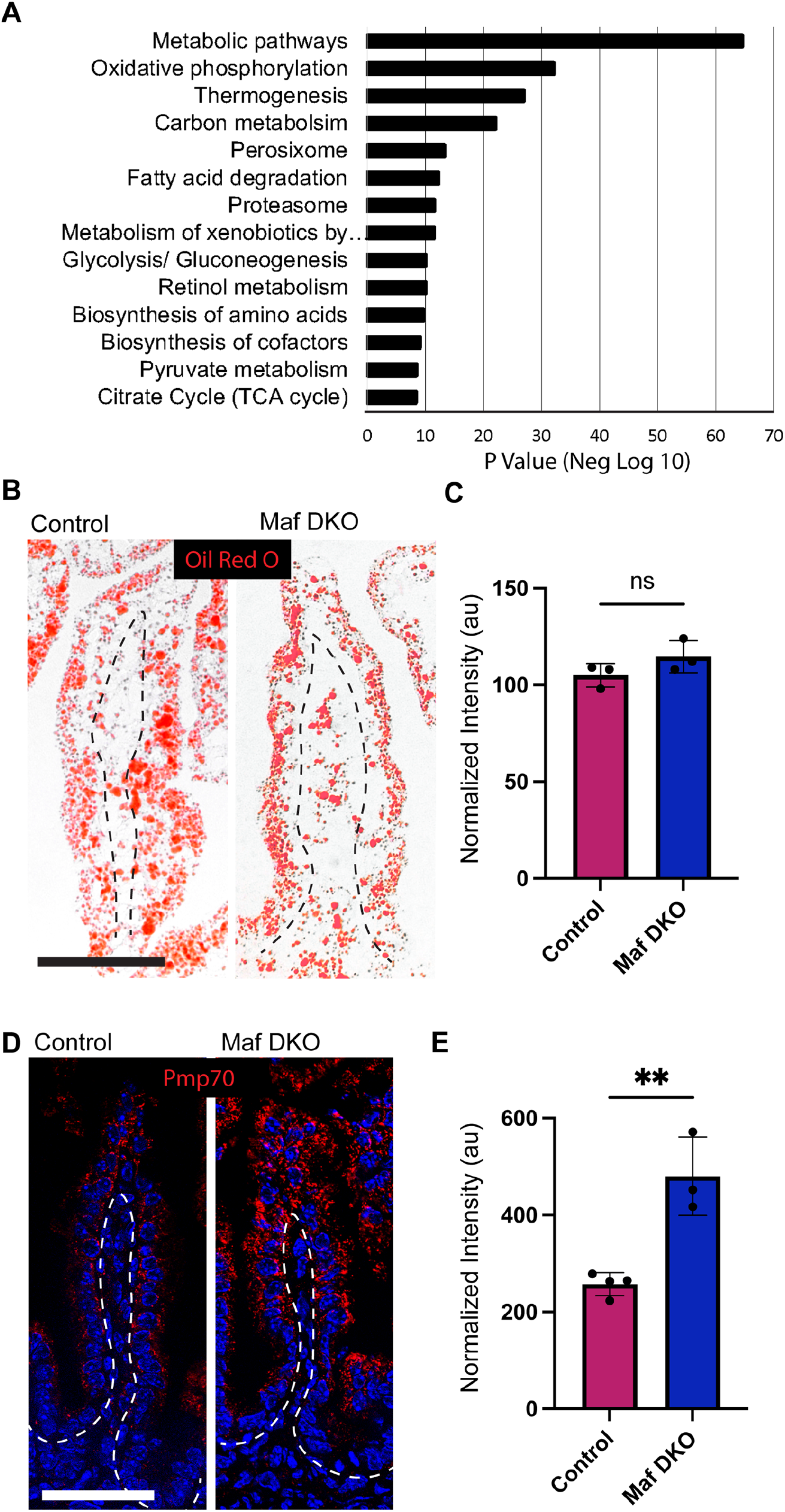
Increased expression of metabolic genes Maf DKO neonates. (A) KEGG analysis of genes up-regulated in Maf DKO villi. Disease terms excluded from list. (B) Oil Red O staining on proximal intestines of mice gavaged with 15 μL corn oil and collected 3h later. (C) Quantification of Oil Red O signal in Maf DKO and control mice. n=3 control and 3 Maf DKO mice. 5 measurements from 4 different images per mouse were quantified. T-test was performed. (D) Staining for the peroxisomal marker, Pmp70 (red) in P7 medial intestine. (E) Quantification of Pmp70 signal in control and Maf DKO mice, n=3 control and 3 Maf DKO mice. 5 measurements from 4 different images per mouse were quantified. T-Test was performed. *dashed lines indicate basement membrane, scale bars are 50 μm.

### Premature expression of adult enterocyte genes in Maf DKO neonates

In addition to metabolic genes for fatty acid degradation, mRNAs for a number of brush border enzymes, including sucrase-isomaltase (Sis) and trehalase (Treh), are increased in Maf DKO neonates (Figure 7D). We validated these changes at the protein level as well. Immunohistochemistry of sucrase-isomaltase revealed increased brush border staining in the Maf DKO intestine, as compared to control littermates (Figure 7A), consistent with its localization in adult enterocytes. Trehalase levels were also increased in the mutant mice by western blot analysis (Figure 7B). Finally, arginase 2 (Arg2), a gene that is important for urea metabolism is also upregulated at the protein level (Figure 7C).

**Figure 7.**
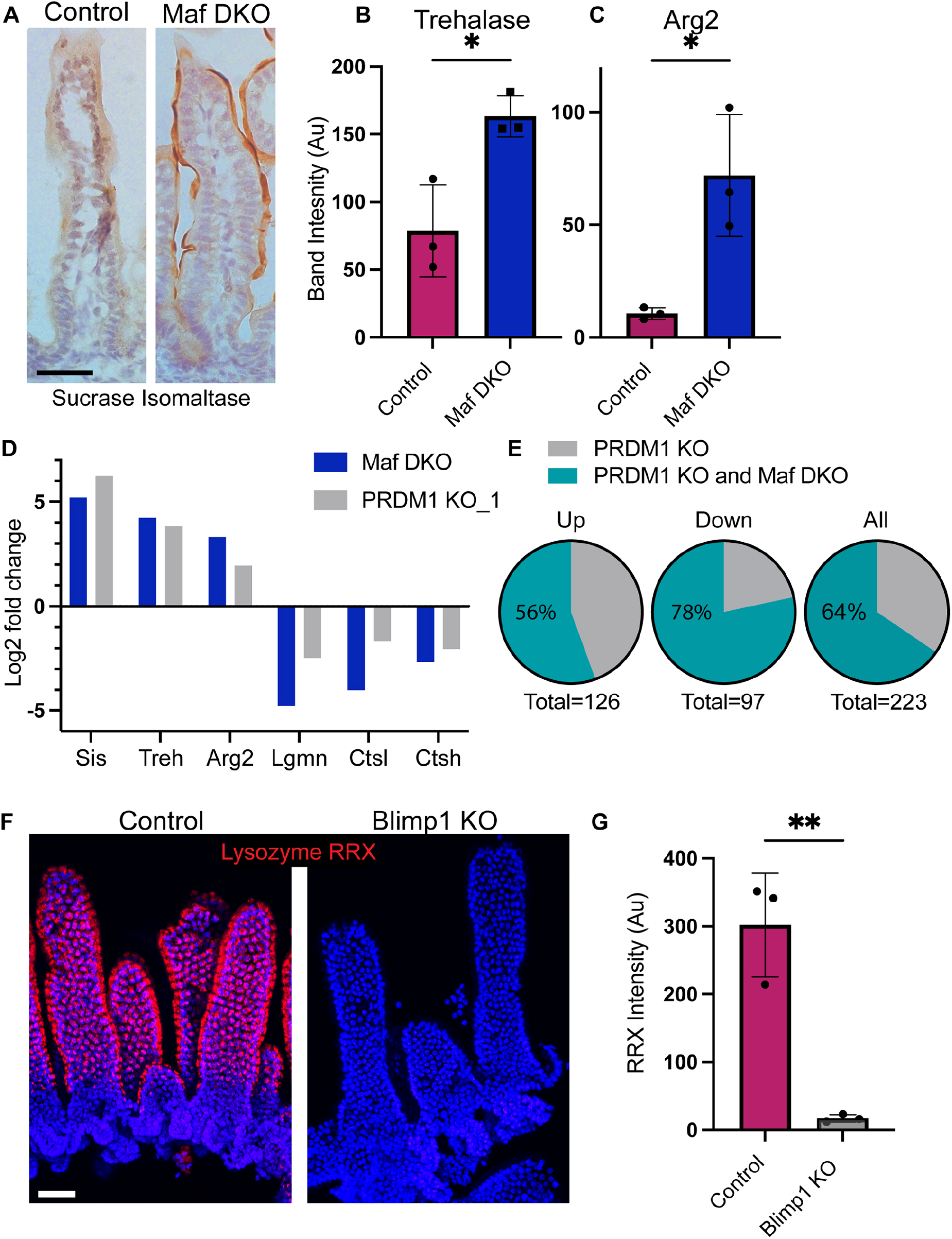
Prdm1 KO neonates display similar phenotype to Maf DKO neonates. (A) Immunohistochemistry analysis of sucrase isomaltase in P7 intestine of control and Maf DKO mice. (B and C) Western blots of protein isolated from distal intestine of P7 mice n=3 control and n=3 Maf DKO showing increased protein expression of Arg2 (p=0.0174) and Trehalase (p= 0.0177) Unpaired T-Test. Band intensity was normalized to tubulin loading control. (D and E) Comparisons of Maf DKO sequencing hits and published array data following Prdm1 KO, on a select set of genes (D) and on the entire gene list (E). (F) Protein uptake assay on Prdm1 KO mice. Oral gavage of 40 μg Lys-RRX (red) into the stomachs of P7 mice, intestines collected 3h later. Epithelial whole mounts are shown. (G) Quantification Lys-RRX brightness in control and Maf DKO. n=3 control and 3 Prdm1 KO, 5 measurements from 4 different images per mouse were quantified. T-Test was run.

Strikingly, many of these genes were also differentially regulated following deletion of Prdm1 in the intestinal epithelium (Harper et al., 2011; Muncan et al., 2011). The Prdm1 gene encodes the Blimp1 transcriptional repressor which is expressed in the intestine from late embryonic development and throughout weaning, and functions to repress the expression of mature enterocyte genes. Loss of Blimp1 leads to a failure to thrive in neonatal mice, and premature maturation of neonatal enterocytes (Harper et al., 2011; Muncan et al., 2011). Sucrase isomaltase, trehalase, and Arg2, which are expressed in mature enterocytes (Henning, 1981; Henning, 1985; Hurwitz and Kretchmer, 1986) are inappropriately expressed in neonatal intestines of Maf DKO and Prdm1/Blimp1 KO mice (Figure 7D). Further, there was a similar decrease in lysosomal protease gene expression in Prdm1/Blimp1 KO and Maf DKO intestines (Figure 7D). These are just a few examples highlighting an overall shift in the expression of Maf DKO enterocytes towards an adult enterocyte signature. Thus, we sought to compare all the differentially regulated genes identified by microarray in P7 Prdm1 KO intestines (Muncan et al., 2011) to those identified as differentially regulated in Maf DKO intestines. We determined that 133 out of 208 genes identified as being significantly changed in Prdm1 intestine by microarray analysis were also differentially regulated in Maf DKO mice (Figure 7E). The hypergeometric test indicates that the enrichment is 4.5 fold over random (p=4.6e-65).

Given these similarities, we next asked whether there were similar deficits in nutrient uptake. Using the same gavage scheme described above, we found that Prdm1/Blimp1 KO enterocytes fail to take in fluorescently labeled protein (Figure 7F, G). These data suggest that these factors may either act together or regulate each other to control nutrient absorption. MafB and cMaf mRNA levels were not significantly altered in the Prdm1 KO microarrays. In contrast, we noted a small decrease (0.58 fold) in Prdm1 mRNA levels in Maf DKO intestines (Figure 7, Supplement 1). However, antibody staining detected similar levels of Blimp1 positive nuclei in Maf DKO intestines (Figure 7, Supplement 1). Together, these data support these proteins acting in concert rather than regulating each other’s expression.

## Discussion

In this work, we demonstrated that Maf family transcription factors, MafB and cMaf, are required for bulk nutrient absorption in the neonatal intestine. These factors, whose expression is controlled by master enterocyte regulators HNF4**α/**γ, are found specifically in enterocytes in the intestinal epithelium from their earliest emergence through adulthood. MafB and cMaf are required for bulk internalization of nutrients, and their loss results in decreased expression of lysosomal, phagocytic, and endosomal genes. Notably, there is also an upregulation of many metabolic pathways, including fatty acid degradation pathways in the peroxisome. Similar functional defects and gene expression changes are also found in Prdm1/Blimp1 mutant intestines. Together, this work defines important transcriptional regulators of neonatal enterocyte function, and identifies both putative machinery for nutrient uptake, and potential compensatory pre-maturation of neonatal enterocytes when bulk intake pathways are perturbed.

While there is clear heterogeneity in enterocyte function both spatially and temporally, the underlying transcriptional regulation of this has remained unclear. HNF4**α/γ** factors are required for a general enterocyte program, and regulate pathways for brush border assembly, and metabolic processes (Chen et al., 2021; Chen et al., 2019a). However, we have limited knowledge of the downstream regulators that control the specialized functions of different types of enterocytes. As an example, Gata4 is expressed in the duodenum and jejunum but not the ileum and loss of Gata4 leads to enterocyte expression that is typically associated with the illeal region of the intestine (Bosse et al., 2006). Our data demonstrate that Maf factors lie downstream of HNF4**α/γ** and strongly suggest that they are direct targets of these transcriptional regulators, based on RNASeq, ChIP-Seq and chromatin conformation analysis. However, while HNF4**α/γ** are expressed throughout the crypt/villus axis (Chen et al., 2019b), Maf factors are only present in enterocytes, suggesting additional levels of regulation to control Maf expression.

Maf expression is first evident as villi begin to form. While many Maf expressing cells on newly-formed embryonic villi begin to lose expression of proliferative marker, Ki67, there are a portion that retain this marker. By the time the mice are born, there is a clear delineation between the postmitotic Maf positive cells of the villi and the proliferative inter-villar cells. Indeed, lineage tracing of MafB-expressing cells demonstrated that their progeny do not significantly contribute to crypts, the stem cell niche. This was surprising given a prior study which suggested that all embryonic intestinal epithelial cells are equally capable of contributing to the stem cell pool. Mechanistically, this model suggested fission of embryonic villi repositions enterocytes into inter-villar regions where they continue to proliferate (Guiu et al., 2019). Our data demonstrate that at least some embryonic epithelial cells (MafB positive cells) are committed to their differentiated fate and do not produce stem cell progeny but do not rule out that MafB negative cells of the emerging villi can contribute to stem cell pools. However, it is important to note that the data supporting this model are based on K19-Cre^ER^ based lineage tracing and the assumption cells throughout the intestinal epithelium are evenly labeled. We found that under the conditions used, recombination preferentially occurs in inter-villar regions, not in an unbiased way throughout the entire epithelium, thus lending support to the model that most cells on embryonic villi do not contribute to the stem cell population.

Though Maf factors are expressed in enterocytes throughout life, our data suggest they are most important for intestinal function in neonates. Maf DKO mice exhibited failure to thrive, with weights only about 70% of their wild type littermates. The Maf mutant intestine, while largely normal histologically, had a profound defect in the uptake of large macromolecules from the lumen. This mode of nutrient absorption is specific to the neonate where endocytic and phagocytic uptake followed by degradation in the lysosome is thought to play a major role (Gonnella and Neutra, 1984; Wilson et al., 1991).

The machinery for bulk uptake in neonatal enterocytes remains largely unstudied. Work in zebrafish, which are thought to use similar uptake pathways throughout their lives, has identified important roles for scavenger receptors, including cubilin and endocytic adaptors, such as Dab2 (Park et al., 2019). The function of Dab2 is conserved in neonatal mice, as we demonstrated lack of efficient protein uptake in Dab2 mutant neonatal intestines (Park et al., 2019). The down-regulated genes in Maf knockout mice provide a potential parts list for genes involved in bulk uptake of nutrients. Notably, we find that many endocytic, phagocytic and lysosomal genes, including Dab2, are down-regulated. In addition to these pathways, many actin regulators were identified in the genes that are both bound by MafB (by CUT&RUN) and down-regulated in the Maf mutant intestine. Of the 13 genes in this category, 11 are actin regulators that have been demonstrated to play important roles in endocytosis/phagocytosis. These endocytic and cytoskeleton genes are excellent candidates for future function validation of roles in nutrient uptake.

In addition to the down-regulated genes in the Maf DKO, there was also a substantial number of genes that were upregulated. These include many “adult” enterocyte genes and suggest that either Mafs inhibit their expression directly, or their upregulation is secondary to disruption of other genes and pathways following loss of Mafs. Loss of bulk uptake of nutrients may lead to increased expression of adult genes to compensate for the lack of nutrient uptake. A very similar set of differentially regulated genes was identified in the Prdm1/Blimp1 mutant intestine. Blimp1 is a transcriptional repressor proposed to inhibit adult enterocyte gene expression in neonates (Harper et al., 2011; Muncan et al., 2011). Remarkably, we find that Prdm1/Blimp1 is also required for bulk nutrient uptake in neonates, consistent with the previously published gene expression data. The nature of the interaction of Maf factors and Blimp1 remains unclear in the intestine, though there is precedent for these factors working cooperatively to induce the expression of IL-10 in CD4+ T cell subsets (Neumann et al., 2014).

Finally, the intriguing finding that many metabolic pathways are upregulated in both Maf and Prdm1 knockout intestine demonstrates a close integration of metabolism and physiology in these cells. Notably, we found that the mice, while deficient in bulk uptake via phagocytosis and endocytosis, were able to take in lipids similar to control littermates. This is likely because, in some forms, lipids do not rely on bulk uptake (Johnston and Borgstroem, 1964; Shiau, 1981). In Maf DKOs, there is upregulation of almost all enzymes of the fatty acid beta-oxidation pathway as well as peroxisomal genes and number of peroxisomes. These data suggest that these mice may become more reliant on energy and/or building blocks from fatty acid catabolism. It is not clear whether many of these changes are directly controlled by these transcription factors and/or whether they are controlled by changes in the nutritional status of the mice due to loss of bulk uptake. Previous work demonstrated that enterocytes change their expression of digestive enzymes in response to the nutrients available. Upon prolonged exposure to a lactose rich diet, enterocytes maintained the expression and activity of lactase (Peuhkuri et al., 1997). Together, these data underscore the dynamic modes of nutrient uptake, the capacity of enterocytes to utilize the macromolecules available, and the transcriptional network that governs these changes.

## Materials and Methods

### Mice

Animal work was performed in accordance with Duke University’s Institutional Animal Care and Use Committee guidelines and approval. Mouse lines used in this study were: CD1 (Charles River), MafB-Cre (Wu et al., 2016), mTmG (Muzumdar et al., 2007), Villin-Cre (Madison et al., 2002), Villin-Cre^ER^ (gift from Sylvie Robine, Institut Curie (el Marjou et al., 2004), MafB^fl/fl^ (Yu et al., 2013), cMaf^fl/fl^ (Wende et al., 2012)), and Prdm1^fl/fl^ (Shapiro-Shelef et al., 2003). Both male and female mice were used and genotyping was performed by PCR. Mice were maintained in a facility with 12h light/dark cycles.

### Tissue Preparation

Isolated intestinal tissue was either embedded immediately in Optical Cutting Temperature (Sakura) or prefixed in 4% paraformaldehyde in PBS-T (PBS + 0.2% Triton X-100) overnight before embedding. Proximal, medial, and distal regions, corresponding to duodenum, jejunum and ileum were collected. The frozen tissue in blocks were cut in 7μm sections using a cryostat. Blocks and tissue sections were stored at −80. For each experiment, a minimum of 3 age matched mice from each condition were collected and analyzed.

### Intestinal epithelium Isolation

Mice were sacrificed and intestines were dissected. Several ∼1cm pieces of small intestine were isolated, cut open longitudinally and washed in PBS. Any luminal contents were removed by gentle shaking. Up to 5, ∼1cm sections were incubated in 5mL of 30mM EDTA in PBS. Intestines from P7 mice were incubated at 37°C for 10 min without rotation and intestines from adult mice were incubated for 30min at 4°C while rotating. The intestines were transferred to a petri dish with PBS, grasped with forceps and then shaken to release the epithelium. The epithelial sheets were transferred to a 15mL conical tube and allowed to gravity pellet. The tissue was further processed according to the downstream application.

### Single cell isolation from intestinal epithelium

Intestinal epithelium was incubated in PBS + 0.8mg/mL dispase for 20min at 37C with intermittent shaking. Cells were then passed through a 70μm strainer, washed with PBS +10% FBS and cells were pelleted for 5min at 2400 RPM. Supernatant was aspirated and cells were washed again with PBS +10% FBS. Cells were either fixed in 4% PFA in PBS for 15min at RT, washed 2x with PBS, and stored at 4C or used fresh.

### Immunofluorescence/IHC

After thawing the tissue sections, they were fixed with 4% PFA in PBS-Triton, with 0.2% TritonX-100, for 8min. When staining for Muc2, antigen retrieval was performed by boiling samples for 10min in 10mM sodium citrate buffer pH 6.0. Tissue sections were washed with PBS-Triton, then blocked with 3% Bovine Serum Albumin (BSA) (Cytiva), 5% Normal Goat Serum (NGS)(Gibco), 5% Normal Donkey Serum (NDS)(Sigma) in PBS-Triton for 15 min. Primary antibody were diluted in blocking buffer and added to the sections for 15min – 1h at RT. After 3x washes with PBST, secondary antibody and any stains, such as Hoescht or Phalloidin, were added to the tissue sections for 15min at RT. After 3x washes with PBST, the slides were mounted with 90% glycerol in PBS plus 2.5 mg/ml p-Phenylenediamine (Sigma-Aldrich) and sealed with clear nail polish. The sections were imaged using Zeiss AxioImager Z1 microscope with Apotome.2 attachment, Plan-APOCHROMAT 20X/0.8 objective or Plan-NEOFLUAR 40X/1.3 oil objective, Plan-APOCHROMAT 63x/1.4oil objective, Axiocam 506 mono camera, and Zen software (Zeiss).

### IHC

After thawing the tissue sections, they were fixed with 4% PFA in PBS-Triton for 8min. When staining for Lamp1 and Sucrase Isomaltase, antigen retrieval was performed by boiling samples for 10min in 10mM NaCitrate buffer pH 6.0. Tissue was blocked first with 0.3% hydrogen peroxide in PBS for 10min, then in 3% BSA, 5% NGS, 5% NDS in PBS-Triton for 15 min at RT. The tissues were incubated with primary antibody diluted 1:100 in blocking solution for 1h at RT then washed 3x with PBS-Triton. Next, the tissues were incubated with HRP secondary antibodies 1:100 dilution in blocking solution for 15min then washed 3x with PBS-Triton. Dab reagent (Vector Labs) was prepared fresh according to the directions in the kit then added to the tissue sections for 30s to 10min. Tissue sections were washed, then allowed to dry completely. Slides were mounted with Permount (Fisher Scientific) and imaged as described above.

### Immunostaining intestinal epithelial whole mounts

After isolating intestinal epithelium as described above, the tissue was fixed overnight in 4% PFA in PBS at 4C. For immunostaining, the tissue was blocked in 3% BSA, 5% NGS, 5% NDS in PBST for 45min while rocking at RT, incubated with primary antibody diluted in blocking solution for 1.5h while rocking at RT. Tissue was allowed to gravity pellet, and washed 3x for 5min while rocking. Secondary antibody along with nuclear stain (Hoescht) and stains for actin (Phalloidin) were diluted in blocking solution and incubated for 45 min at RT while rocking and protected from light. In cases where only stains or nuclear dye were used, the blocking and primary steps were not performed. Epithelial whole mounts were mounted on slides using melted VALAP (vasaline, lanolin, and paraffin in a 1:1:1 ratio) to create a border in which the tissue was placed. After the VALAP had solidified, the tissue was pipetted inside the border and excess liquid was wicked away. 90% glycerol in PBS plus 2.5 mg/ml p-Phenylenediamine (Sigma-Aldrich) was added to cover the tissue and a coverslip was placed on top and sealed with melted VALAP. Slides were stored at 4C and imaged within 1wk of preparation on Zeiss 780 upright confocal with a 20X/0.8 Plan-Apochromat objective or 63X/1.4 oil immersion Plan-Apochromat objective.

### Western Blot

After isolating intestinal epithelium as described above, 4X, by volume, of sample buffer with 15% BME, was added to the tissue which was then incubated at 95C for 10 minutes with intermittent trituration. Samples were aliquoted, flash frozen, and stored at −80C. Samples were run out on 10% polyacrylamide gels, transferred to nitrocellulose membranes. Membranes were blocked in 5% BSA in PBS-Tween for 45min at RT. Primary antibodies were diluted in 5% BSA in PBS-Tween (1% Tween) and added to the membrane overnight at 4C while rocking. Tubulin is the exception to this and was added to the membranes for 30min at room temperature while rocking. Membranes were washed 3x for 5min with PBS-Tween. LI-COR secondary antibodies were diluted in 5% BSA in PBS-Tween and added to the membrane for 45min at RT while rocking. After washing, membranes were imaged using the LI-COR Odyssey and band intensity was quantified using LI-COR software. Protein amounts were standardized using Revert total protein stain (LI-COR). Additionally, protein levels were normalized to tubulin for quantification.

### Oil Red O Stain

P7 mice were gavages with 15μL corn oil. 3h later, intestines were collected as described above. This Oil Red O stain protocol is based on the one found at ICHworld.com. A stock solution of 0.5g Oil Red O (Sigma) in 100mL isopropanol was prepared. Working solution was prepared fresh by adding 15mL of stock solution to 10mL water, incubating at RT for 10min then gravity filtering through Whatman paper. At the same time, slides with 7μm thick small intestine sections were dried completely then fixed with 4% PFA in PBS for 1h at RT. Slides were washed 3x with diH_2_O then incubated in the filtered Oil Red O working solutions for 30min at RT. Slides were washed 3x again and run under tap water for 5mins. Slides were dried completely then mounted with Permount.

### In Vivo Uptake Assays

P7 mice were gavaged with 40μg Lysozyme-RRX (Lys_RRX), 40μg Dextran-GFP (Dex-488) in a total volume of 10μL, or 15μL corn oil using 22 gauge plastic feeding tubes. Mice were euthanized and intestines were collected 3h post gavage. Pieces of whole intestine, epithelial isolates, and single cells were collected as described above depending on the experiment.

### FACS analysis

Fixed intestinal epithelial cells were isolated as descried above and stored at 4C until the time of analysis. Immediately before FACS analysis, the cells were passed though 100μm Celltrix filters. Samples were processed on Fortessa Analyzer with assistance from the Flow cytometry core at Duke. Samples were normalized to a no gavage control.

### RNA sequencing

Epithelial cells were isolated from proximal, medial and distal P7 intestine of 3 control and 3 Maf DKO mice as described above and pooled into a single tube per mouse. RNA was isolated using the Qiagen RNAeasy kit. The RNA concentration for each sample was determined via nano drop, then samples were aliquoted and flash frozen. Samples were sent to Novogene for sequencing and analysis. Genes with changes greater than 0.2 log2 fold change or less than −0.25 log fold change and had a P value of less than 0.05 were considered differentially regulated. Raw data has been submitted to GEO (accession number pending).

### CUT&RUN

Cells were isolated from the distal intestine of a P7 CD1 mouse. Intestinal epithelial cells were isolated and further processed into single cells as described above. The CUT&RUN Assay Kit from Cell Signaling Technologies was used to perform the CUT&RUN. The MinElute PRC Purification Kit from Quiagen was used to isolate the DNA. Libraries were prepared as previously described (Meers et al., 2019; Skene and Henikoff, 2017). The concentration of the libraries was determined via Qubit and equal amounts of sample were pooled and sent to Novogene for sequencing. In brief the results were analyzed by standard pipeline: trimgalore -> bowtie2 alignment -> calling peaks by MACS2. Raw data has been submitted to GEO (accession number pending).

### Statistical Analysis

Figure legends contain details about stats for each figure. Generally, T-tests were performed using PRISM software and p values were reported as follows: ns = not significant’ *, p < 0.05; **, p < 0.01, ***, p < 0.001.

| REAGENT or RESOURCE | SOURCE | IDENTIFIER |
| --- | --- | --- |
| **Antibodies** |  |  |
| MafB: Dilution used for IF: (1:250) | Novus | NBP1-81342<br>RRID: AB_11005476 |
| cMaf: Dilution used for IF: (1:100) | Invitrogen | PA5-23179<br>RRID: AB_2540705 |
| Ki67: Dilution used for IF : (1:100) | Invitrogen | 14-5698-80<br>RRID: AB_10853185 |
| Muc2: Dilution used: (1:100) | Novus | NBP1-31231<br>RRID: AB_10003763 |
| Lamp1: Dilution used for IF: (1:100), for WB (1:1000) | Abcam | ab24170<br>RRID: AB_775978 |
| Gulp: Dilution used for WB: (1:1000) | Millipore Sigma | HPA020587<br>RRID: AB_1850452 |
| Legumain: Dilution used for WB: (1:1000) | ABclonal | A6829<br>RRID: AB_2863540 |
| Cathepsin: Dilution used for WB: (1:1000) | ABclonal | A4986<br>RRID: AB_2863404 |
| Pmp70: Dilution used for IF: (1:100) | Abcam | ab3421<br>RRID: AB_2219901 |
| Sucrase Isomaltase: Dilution used for IF: (1:100) | NSJ Bioreagents | RQ4659 |
| Trehalase: Dilution used for WB: (1:1000) | Thermo Scientific | BS-16712R |
| Arg2: Dilution used for WB: (1:1000) | Abcam | ab264066 |
| Blimp1: Dilution used for IF: (1:100) | Santa Cruz | sc-47732,<br>RRID:AB_628168 |
| **Chemicals, peptides, and recombinant proteins** |  |  |
| Phalloidin-TRITC | Sigma | P1591 |
| Phalloidin-488 | Invitrogen | A12379 |
| Lysozyme_RRX | Nanocs | LS1-RB-1 |
| Dextran GFP | Invitrogen | D22910 |
| mCherry | Gift from Michel Bagnat |  |
| **Commercial assays** |  |  |
| DAB (3,3'-diaminobenzidine) Peroxidase (HRP) Substrate Kit, With Nickel | Vector Laboratories | SK-4100 |
| RNeasy Kit | Qiagen | 74104 |
| MinElute PCR Purification Kit | Qiagen | 28004 |
| **Experimental models: Organisms/strains** |  |  |
| Mouse: CD1 IGS | Charles River | Strain Code: 022 |
| Mouse: MafB-Cre | The Jackson Laboratory | Strain Code: 029664 |
| Mouse: RGT | The Jackson Laboratory | Strain Code: 007576 |
| Mouse: K19-Cre | The Jackson Laboratory | Strain Code: 026925 |
| Mouse: Villin-Cre | The Jackson Laboratory | Strain Code: 021504 |
| Mouse: Villin-CreER | The Jackson Laboratory | Strain Code: 020282 |
| Mouse: MafB<sup>fl/fl</sup> | The Goodrich Lab | MGI:5581666 |
| Mouse: cMaf<sup>fl/fl</sup> | The Birchmeier Lab | MGI:5316775 |
| Mouse: Prdm1<sup>fl/fl</sup> | The Jackson Laboratory | Strain Code: 008100 |

## Acknowledgements

We thank members of the Lechler Lab for comments on the manuscript, and Carmen Birchmeier, Lisa Goodrich, and Michel Bagnat for reagents. In addition, we thank Bin Li from the Duke Flow Cytometry Shared Resource for cell sorting assistance, Yasheng Gao from the Duke Light Microscopy Core facility for imaging assistance, and Jianhong Ou for CUT&RUN analysis. Grant support: R01DK121915 and R01DK126446 (to M.Z.), R01DK11798 and R01AR067203 (to T.L.) and R35HG011328 and U01HL156064 (to Y.D.).

---

**Figure 1, Supplement 1.**
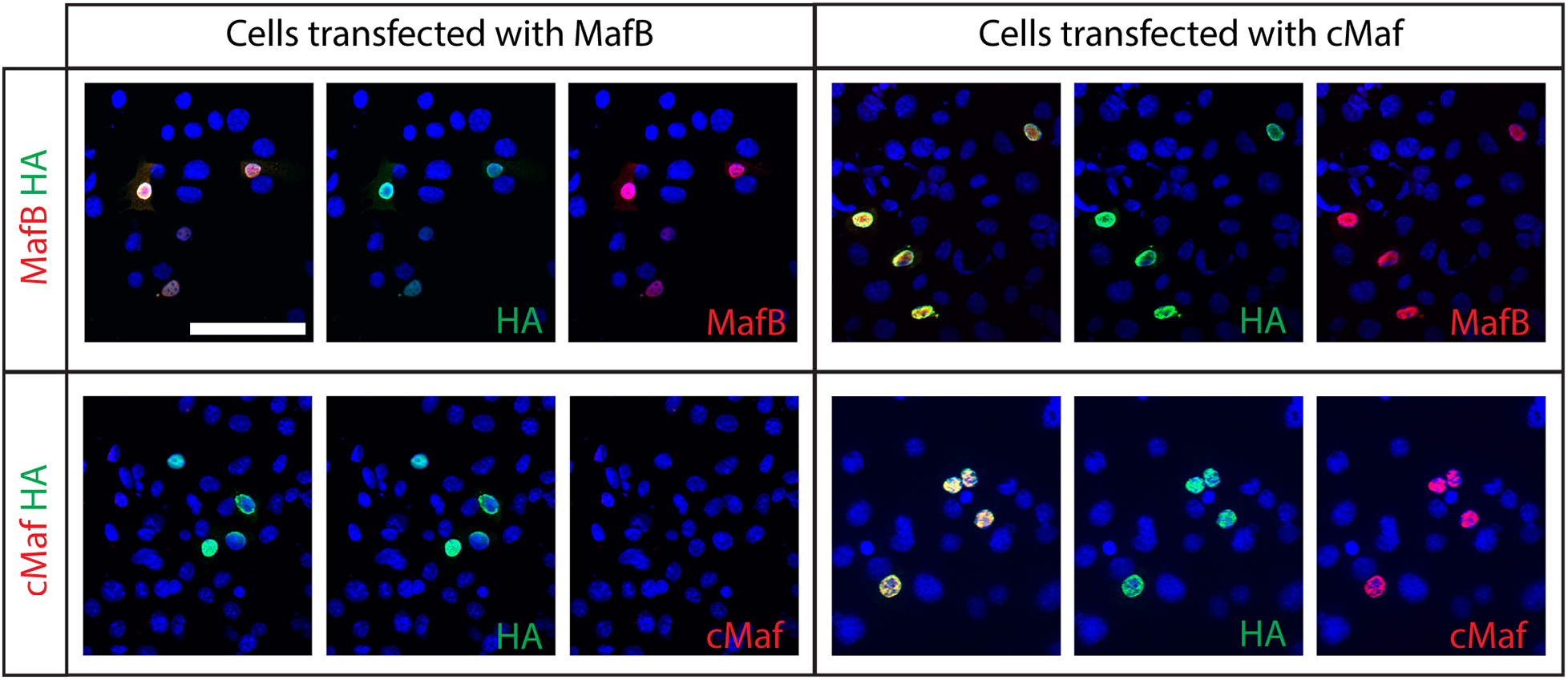
Maf antibody specificity. Keratinocytes were transfected with plasmids expressing either MafB-HA (left) or cMaf-HA (right) for 24h, fixed, then stained with HA and MafB (top) or cMaf (bottom) antibodies.

**Figure 1, Supplement 2.**
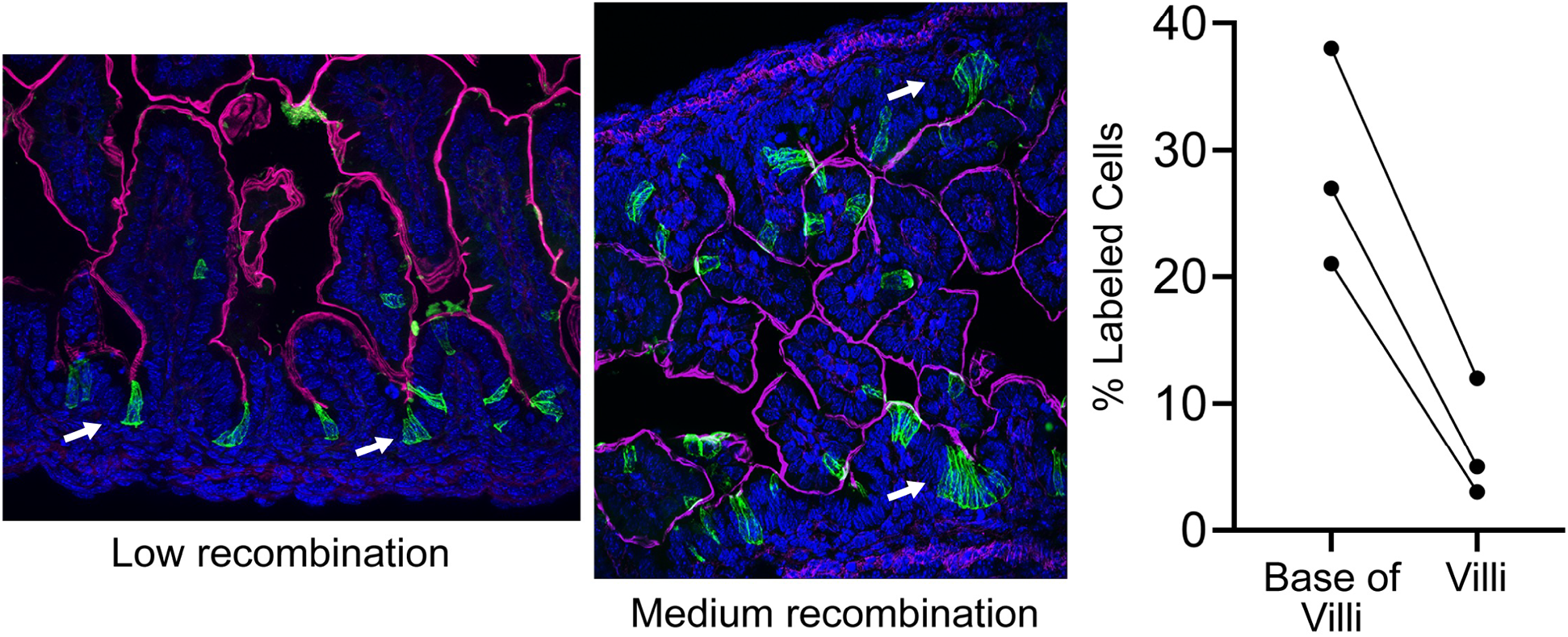
K19-Cre^ER^ predominantly labels the inter-villar proliferative cells of the embryonic intestine. Timed mating with K19-Cre^ER^ x mTmG mice. Dam was gavaged with tamoxifen at E16.5, intestines were isolated from embryos at E17.5. Recombined cells (green), actin (red), arrows indicate examples of GFP lineage labeled cells in inter-villar spaces. While levels of recombination varied between mice, low recombination (left) and medium recombination (right), the percent of labeled cells in the inter-villar spaces was higher than the percent labeled villar cells. n=3 mice.

**Figure 3, Supplement 1.**
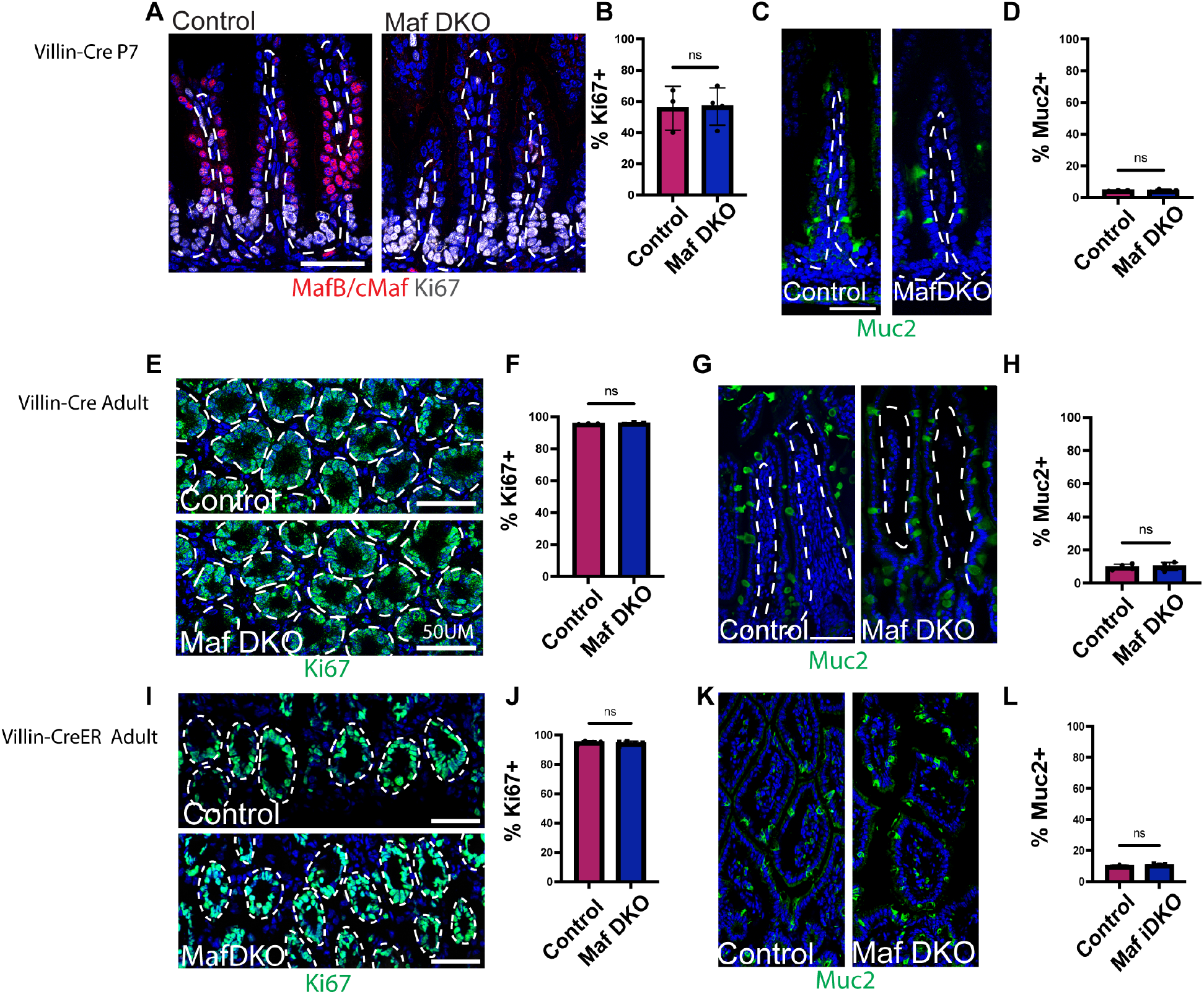
Loss of Maf proteins affects neither proliferation nor lineage specification. (A) Ki67 (white) in P7 control and Maf DKO mice. (B) Quantification of the percent of Ki67+ cells in the inter-villar spaces (n=3 control mice, cells counted per mouse: 173, 207, 114. Maf DKO mice: N=4 mice, cells counted per mouse: 120, 169, 149 and 206.) (C) Muc2 (green) in P7 control and Maf DKO mice. (D) Quantification of the percent of Muc2+ cells (n= 3 control mice, cells counted per mouse:557, 668, 455. n=4 Maf DKO mice,cells counter per mouse: 722, 711, 626, 606.) (E) Ki67 (green) in adult control and Maf DKO mice. (F) Quantification of percent Ki67+ cells in crypts (n=3 control mice, cells counted per mouse: 390, 471, 504. n=3 Maf DKO mice, cells counted per mouse: 427, 427, 390.) (G) Muc2 (green) in adult control and Maf DKO mice H. Quantification of percent of Muc2+ cells (n=4 control mice, cells counted per mouse: 1065, 859, 786. n=4 Maf DKO mice, cells counted per mouse: 1064, 1036, 793, 999.) (I) Ki67 (green) in adult control and Maf iDKO. (J) Quantification of percent Ki67+ cell in crypts (n= 4 control mice, cells counted per mouse: 506, 490, 488, 792. n=5 Maf iDKO, cells counted per mouse: 601, 723, 1174, 801, 841. (K) Muc2 (green) in adult control and Maf iDKO (L) Quantification of percent Muc2+ cells. n=3 control mice, cells counted per mouse: 2693, 3735, 1382. n=5 Maf iDKO, 2885, 3335, 1985, 2046, 1398 White dashed lines indicate basement membrane, scale bars are 50 μm.

**Figure 4, Supplement 1.**
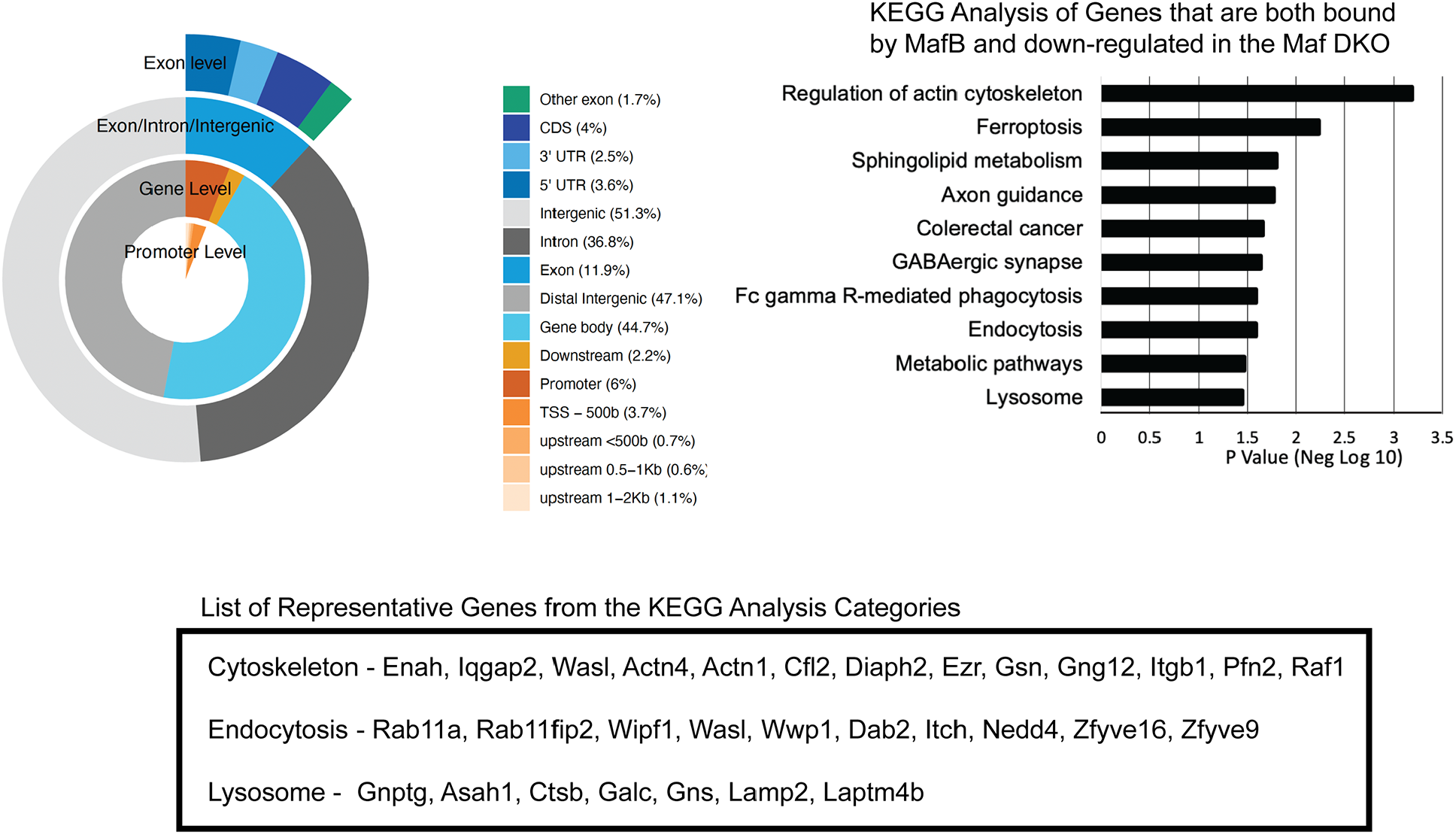
CUT&RUN analysis of MafB in the intestinal epithelium. Image on left shows analysis of binding sites on chromatin. KEGG analysis of genes that are both bound by MafB and down-regulated in the Maf DKO intestine are shown on right, with individual genes in relevant categories listed below.

**Figure 7, Supplement 1.**
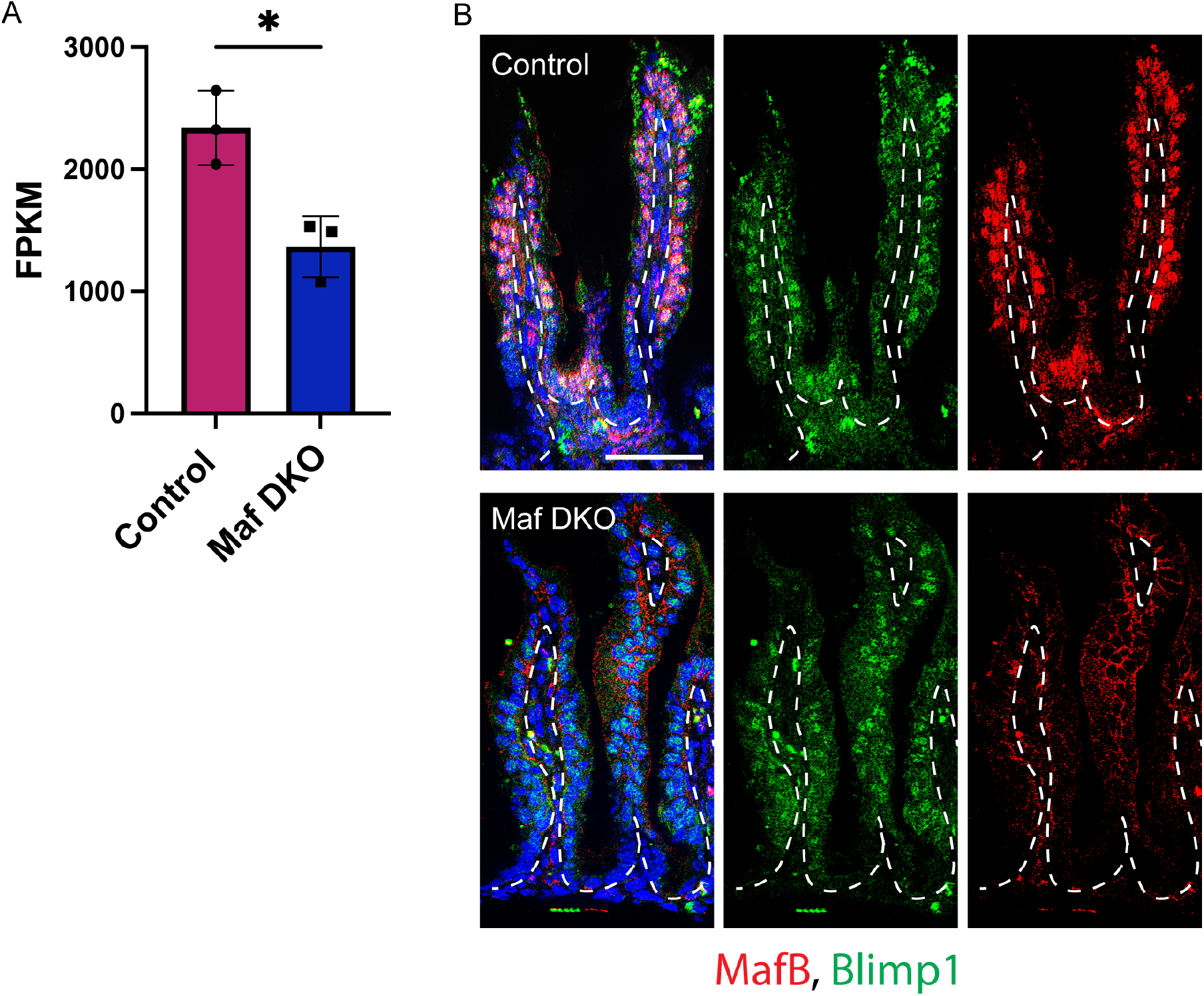
Prdm1/Blimp1 expression in Maf DKO intestines. (A) RNA seq of Maf DKO intestinal epithelium, Prdm1 gene changes. (B) Immunofluorescent images of P7 medial intestine stained for MafB (red) and Blimp1 (green).

## References

Aziz, A., E. Soucie, S. Sarrazin, and M.H. Sieweke. 2009. MafB/c-Maf deficiency enables self-renewal of differentiated functional macrophages. Science. 326:867–871.

Beuling, E., N.Y. Baffour-Awuah, K.A. Stapleton, B.E. Aronson, T.K. Noah, N.F. Shroyer, S.A. Duncan, J.C. Fleet, and S.D. Krasinski. 2011. GATA factors regulate proliferation, differentiation, and gene expression in small intestine of mature mice. Gastroenterology. 140:1219–1229 e1211-1212.

Bosse, T., C.M. Piaseckyj, E. Burghard, J.J. Fialkovich, S. Rajagopal, W.T. Pu, and S.D. Krasinski. 2006. Gata4 is essential for the maintenance of jejunal-ileal identities in the adult mouse small intestine. Mol Cell Biol. 26:9060–9070.

Chen, L., S. Luo, A. Dupre, R.P. Vasoya, A. Parthasarathy, R. Aita, R. Malhotra, J. Hur, N.H. Toke, E. Chiles, M. Yang, W. Cao, J. Flores, C.E. Ellison, N. Gao, A. Sahota, X. Su, E.M. Bonder, and M.P. Verzi. 2021. The nuclear receptor HNF4 drives a brush border gene program conserved across murine intestine, kidney, and embryonic yolk sac. Nat Commun. 12:2886.

Chen, L., N.H. Toke, S. Luo, R.P. Vasoya, R. Aita, A. Parthasarathy, Y.H. Tsai, J.R. Spence, and M.P. Verzi. 2019a. HNF4 factors control chromatin accessibility and are redundantly required for maturation of the fetal intestine. Development. 146.

Chen, L., N.H. Toke, S. Luo, R.P. Vasoya, R.L. Fullem, A. Parthasarathy, A.O. Perekatt, and M.P. Verzi. 2019b. A reinforcing HNF4-SMAD4 feed-forward module stabilizes enterocyte identity. Nat Genet. 51:777–785.

el Marjou, F., K.P. Janssen, B.H. Chang, M. Li, V. Hindie, L. Chan, D. Louvard, P. Chambon, D. Metzger, and S. Robine. 2004. Tissue-specific and inducible Cre-mediated recombination in the gut epithelium. Genesis. 39:186–193.

Gonnella, P.A., and M.R. Neutra. 1984. Membrane-bound and fluid-phase macromolecules enter separate prelysosomal compartments in absorptive cells of suckling rat ileum. J Cell Biol. 99:909–917.

Greengard, O. 1977. Enzymic differentiation of human liver: comparison with the rat model. Pediatr Res. 11:669–676.

Guiu, J., E. Hannezo, S. Yui, S. Demharter, S. Ulyanchenko, M. Maimets, A. Jorgensen, S. Perlman, L. Lundvall, L.S. Mamsen, A. Larsen, R.H. Olesen, C.Y. Andersen, L.L. Thuesen, K.J. Hare, T.H. Pers, K. Khodosevich, B.D. Simons, and K.B. Jensen. 2019. Tracing the origin of adult intestinal stem cells. Nature. 570:107–111.

Haber, A.L., M. Biton, N. Rogel, R.H. Herbst, K. Shekhar, C. Smillie, G. Burgin, T.M. Delorey, M.R. Howitt, Y. Katz, I. Tirosh, S. Beyaz, D. Dionne, M. Zhang, R. Raychowdhury, W.S. Garrett, O. Rozenblatt-Rosen, H.N. Shi, O. Yilmaz, R.J. Xavier, and A. Regev. 2017. A single-cell survey of the small intestinal epithelium. Nature. 551:333–339.

Harper, J., A. Mould, R.M. Andrews, E.K. Bikoff, and E.J. Robertson. 2011. The transcriptional repressor Blimp1/Prdm1 regulates postnatal reprogramming of intestinal enterocytes. Proc Natl Acad Sci U S A. 108:10585–10590.

Henning, S.J. 1981. Postnatal development: coordination of feeding, digestion, and metabolism. Am J Physiol. 241:G199–214.

Henning, S.J. 1985. Ontogeny of enzymes in the small intestine. Annu Rev Physiol. 47:231–245.

Hurwitz, R., and N. Kretchmer. 1986. Development of arginine-synthesizing enzymes in mouse intestine. Am J Physiol. 251:G103–110.

Johnston, J.M., and B. Borgstroem. 1964. The Intestinal Absorption and Metabolism of Micellar Solutions of Lipids. Biochim Biophys Acta. 84:412–423.

Lazarow, P.B., and C. De Duve. 1976. A fatty acyl-CoA oxidizing system in rat liver peroxisomes; enhancement by clofibrate, a hypolipidemic drug. Proc Natl Acad Sci U S A. 73:2043–2046.

Lopez-Pajares, V., K. Qu, J. Zhang, D.E. Webster, B.C. Barajas, Z. Siprashvili, B.J. Zarnegar, L.D. Boxer, E.J. Rios, S. Tao, M. Kretz, and P.A. Khavari. 2015. A LncRNA-MAF:MAFB transcription factor network regulates epidermal differentiation. Dev Cell. 32:693–706.

Madison, B.B., L. Dunbar, X.T. Qiao, K. Braunstein, E. Braunstein, and D.L. Gumucio. 2002. Cis elements of the villin gene control expression in restricted domains of the vertical (crypt) and horizontal (duodenum, cecum) axes of the intestine. J Biol Chem. 277:33275–33283.

Mascre, G., S. Dekoninck, B. Drogat, K.K. Youssef, S. Brohee, P.A. Sotiropoulou, B.D. Simons, and C. Blanpain. 2012. Distinct contribution of stem and progenitor cells to epidermal maintenance. Nature. 489:257–262.

Meers, M.P., T.D. Bryson, J.G. Henikoff, and S. Henikoff. 2019. Improved CUT&RUN chromatin profiling tools. Elife. 8.

Moor, A.E., Y. Harnik, S. Ben-Moshe, E.E. Massasa, M. Rozenberg, R. Eilam, K. Bahar Halpern, and S. Itzkovitz. 2018. Spatial Reconstruction of Single Enterocytes Uncovers Broad Zonation along the Intestinal Villus Axis. Cell. 175:1156–1167 e1115.

Muncan, V., J. Heijmans, S.D. Krasinski, N.V. Buller, M.E. Wildenberg, S. Meisner, M. Radonjic, K.A. Stapleton, W.H. Lamers, I. Biemond, M.A. van den Bergh Weerman, D. O’Carroll, J.C. Hardwick, D.W. Hommes, and G.R. van den Brink. 2011. Blimp1 regulates the transition of neonatal to adult intestinal epithelium. Nat Commun. 2:452.

Muzumdar, M.D., B. Tasic, K. Miyamichi, L. Li, and L. Luo. 2007. A global double-fluorescent Cre reporter mouse. Genesis. 45:593–605.

Neumann, C., F. Heinrich, K. Neumann, V. Junghans, M.F. Mashreghi, J. Ahlers, M. Janke, C. Rudolph, N. Mockel-Tenbrinck, A.A. Kuhl, M.M. Heimesaat, C. Esser, S.H. Im, A. Radbruch, S. Rutz, and A. Scheffold. 2014. Role of Blimp-1 in programing Th effector cells into IL-10 producers. J Exp Med. 211:1807–1819.

Park, J., D.S. Levic, K.D. Sumigray, J. Bagwell, O. Eroglu, C.L. Block, C. Eroglu, R. Barry, C.R. Lickwar, J.F. Rawls, S.A. Watts, T. Lechler, and M. Bagnat. 2019. Lysosome-Rich Enterocytes Mediate Protein Absorption in the Vertebrate Gut. Dev Cell. 51:7–20 e26.

Peuhkuri, K., M. Hukkanen, R. Beale, J.M. Polak, H. Vapaatalo, and R. Korpela. 1997. Age and continuous lactose challenge modify lactase protein expression and enzyme activity in gut epithelium in the rat. J Physiol Pharmacol. 48:719–729.

Robberecht, P., M. Deschodt-Lanckman, J. Camus, J. Bruylands, and J. Christophe. 1971. Rat pancreatic hydrolases from birth to weaning and dietary adaptation after weaning. Am J Physiol. 221:376–381.

Sender, R., and R. Milo. 2021. The distribution of cellular turnover in the human body. Nat Med. 27:45–48.

Shapiro-Shelef, M., K.I. Lin, L.J. McHeyzer-Williams, J. Liao, M.G. McHeyzer-Williams, and K. Calame. 2003. Blimp-1 is required for the formation of immunoglobulin secreting plasma cells and pre-plasma memory B cells. Immunity. 19:607–620.

Shiau, Y.F. 1981. Mechanisms of intestinal fat absorption. Am J Physiol. 240:G1–9.

Skene, P.J., and S. Henikoff. 2017. An efficient targeted nuclease strategy for high-resolution mapping of DNA binding sites. Elife. 6.

Skrzypek, T., J.L. Valverde Piedra, H. Skrzypek, W. Kazimierczak, M. Biernat, and R. Zabielski. 2007. Gradual disappearance of vacuolated enterocytes in the small intestine of neonatal piglets. J Physiol Pharmacol. 58 Suppl 3:87–95.

Sumigray, K.D., M. Terwilliger, and T. Lechler. 2018. Morphogenesis and Compartmentalization of the Intestinal Crypt. Dev Cell. 45:183–197 e185.

Verzi, M.P., H. Shin, L.L. Ho, X.S. Liu, and R.A. Shivdasani. 2011. Essential and redundant functions of caudal family proteins in activating adult intestinal genes. Mol Cell Biol. 31:2026–2039.

Wang, S., C. Cebrian, S. Schnell, and D.L. Gumucio. 2018. Radial WNT5A-Guided Post-mitotic Filopodial Pathfinding Is Critical for Midgut Tube Elongation. Dev Cell. 46:173–188 e173.

Wende, H., S.G. Lechner, C. Cheret, S. Bourane, M.E. Kolanczyk, A. Pattyn, K. Reuter, F.L. Munier, P. Carroll, G.R. Lewin, and C. Birchmeier. 2012. The transcription factor c-Maf controls touch receptor development and function. Science. 335:1373–1376.

Wilson, J.M., J.A. Whitney, and M.R. Neutra. 1991. Biogenesis of the apical endosome-lysosome complex during differentiation of absorptive epithelial cells in rat ileum. J Cell Sci. 100 (Pt 1):133–143.

Wu, X., C.G. Briseno, V. Durai, J.C. Albring, M. Haldar, P. Bagadia, K.W. Kim, G.J. Randolph, T.L. Murphy, and K.M. Murphy. 2016. Mafb lineage tracing to distinguish macrophages from other immune lineages reveals dual identity of Langerhans cells. J Exp Med. 213:2553–2565.

Yu, W.M., J.M. Appler, Y.H. Kim, A.M. Nishitani, J.R. Holt, and L.V. Goodrich. 2013. A Gata3-Mafb transcriptional network directs post-synaptic differentiation in synapses specialized for hearing. Elife. 2:e01341.

Zhang, W., S. Zhang, C. Chen, N. Liu, D. Yang, P. Wang, and F. Ren. 2022. The internalization mechanisms and trafficking of the pea albumin in Caco-2 cells. Int J Biol Macromol.

